# Importin-β specific nuclear transport defects recapitulate phenotypic and transcriptional alterations seen in neurodegeneration

**DOI:** 10.1101/2024.02.14.580338

**Authors:** Jonathan Plessis-Belair, Kathryn Ravano, Ellen Han, Aubrey Janniello, Catalina Molina, Roger Sher

## Abstract

Defects in Nucleocytoplasmic transport have been implicated as an important neurodegenerative pathway in ALS/FTD. Here, we show that a *Nemf*^R86S^ mutation results in the disruption of NCT both *in vitro* and *in vivo*. These disruptions are specific to Importin-β nuclear import, and result in the nuclear loss and cytoplasmic gain of NEMF, Importin-β, and TDP-43. We show that a transient nuclear import block is capable of inducing the mis-localization of TDP-43 and is associated with altered transcriptional expression of ALS, FTD, and AD/ARD genes. Taken together, these findings show that disrupted Importin-β nuclear import, whether through genetic forms such as *Nemf* mutations, or through pharmacological inhibition, is the primary driver of TDP-43 pathology, disease-related transcriptional alterations, and neurodegeneration.

## Introduction

Nucleocytoplasmic transport (NCT) defects underlie several neurodegenerative disorders, including amyotrophic lateral sclerosis (ALS), frontal temporal dementia and frontal temporal lobar degeneration (FTD\FTLD), and Alzheimer’s and Alzheimer’s Related Dementias (AD/ADRD) ^1–3^. Here we show that mutations in Nuclear Export Mediator Factor (NEMF) in mice result in Importin-β-specific nuclear import defects, cytoplasmic protein mis-localization and neurodegeneration, and recapitulate phenotypic and transcriptional alterations found in these neurodegenerative disorders.

NCT involves the regulated trafficking of proteins and RNA between the nucleus and the cytoplasm through the nuclear pore complex (NPC) ^4^. Smaller molecules (<40kDa) can freely migrate through the nuclear pore, whereas larger molecules require active transport mediated by nuclear transport receptors (NTRs, also known as karyopherins) such as importins and exportins ^5^. During nuclear import, nuclear import receptors (NIRs), such as Importin-β, directly bind their cargo in the cytoplasm or utilize importin-α as an adaptor to bind cargo proteins bearing a classical nuclear localization signal (cNLS) ^6,7^. Other NIRs, such as Transportin-1, recognize different nuclear localization signals, such as a Proline-Tyrosine NLS (PY-NLS), for transport into the nucleus ^8^.

Pathological disruption of NCT, such as the mis-localization of Nuclear Pore Complex proteins (Nups), NTRs, and Ran-GTPase, are seen in both animal models and in familial and sporadic forms of ALS, FTD, AD/ADRD ^9–15^. In addition, pathological cytoplasmic protein aggregates mark the progression of most neurodegenerative diseases ^16,17^. One of the most frequently found protein aggregates comprises TAR-DNA binding protein-43 (TDP-43), normally a nuclear protein that, in disease, mis-localizes and aggregates in the cytoplasm ^18^. Cytoplasmic aggregation and loss of nuclear TDP-43 are observed in post-mortem neurons and glia in 97% of ALS, 40% of FTD, and in many cases of AD/ADRD ^19^.

Amino acid substitutions in NEMF have been identified in multiple human patients exhibiting severe neurodevelopmental disorders ^20–22^. However, the mechanisms through which these NEMF variants result in disease are unknown. While NEMF was initially associated with nuclear transport in *Drosophila* (<u>Caliban</u>) ^23^, its canonical role has more recently been elucidated in the context of ribosome quality control (RQC), where it plays a crucial role in targeting partially translated nascent chain polypeptides (NCPs) on the 60S ribosome for proteasomal degradation ^23–28^. NEMF’s interaction with the 60S ribosome facilitates the recruitment and stabilization of the E3 ubiquitin Ligase Listerin (LTN1), promoting the ubiquitination of NCPs ^24,25,27,29–31^. Additionally, NEMF can generate c-terminal alanine (and threonine) tails on NCPs to expose amino acids in the ribosomal exit tunnel for subsequent ubiquitination by Listerin ^30,32,33^. Mutations in both *Listerin* and *Nemf* in mice have been associated with neurodegenerative diseases ^20,34^. Notably, Importin-β has been demonstrated to bind to translating NCPs targeted to the nucleus, suggesting that protein quality control and nuclear import mechanisms are more directly interconnected ^35^.

In this study, we have demonstrated that the mutant mouse model, *Nemf^R86S^*, exhibits impaired nuclear import of proteins containing canonical nuclear localization signals (cNLS), consequently leading to defects in nucleocytoplasmic transport both *in vitro* and *in vivo*. We have identified a specific defect in Importin-β mediated transport that manifests as both nuclear loss and cytoplasmic gain of NEMF, Importin-β, and TDP-43, along with a collapse of the Ran gradient. This dysregulation is further associated with the altered expression of ALS-linked genes, specifically *Stmn2*. Remarkably, the dysregulation of ALS-linked genes observed in our *Nemf* mutant-mouse model can be reproduced through the transient induction of a nuclear import block in both murine and human cells using a small molecular antagonist of Importin-β, importazole. Our study reveals that there is a causative role for an Importin-β nuclear import deficiency upstream of the multiple protein and transcriptional pathologies found in neurodegenerative disease.

## Results

### Motor Neuron Degeneration is associated with Progressive Nuclear loss of mutant NEMF in Lumbar Spinal Cord

A previous study demonstrated that mutant *Nemf*^R86S^ mice exhibit progressive neuromuscular degeneration, characterized by a loss of neuromuscular junctions (NMJs) and a progressive axonal degeneration ^20^. To investigate the mechanism by which the missense *Nemf* mutation led to an age-dependent neurodegenerative phenotype associated with motor deficits, we immunostained transverse lumbar spinal cords of wild type and *Nemf*^R86S^ homozygous mutant mice. Lumbar spinal motor neurons were identified by ChAT-positive cells located in the ventral horn. In wild type (WT) mice, NEMF was found to be predominantly nuclear with diffuse cytoplasmic staining (Fig A, F). *Nemf*^R86S^ mice exhibited nuclear loss of NEMF with a significant cytoplasmic accumulation in the somata of ChAT positive motor neurons (Fig 1A,F). However, this phenotypic pathology is not exclusive to spinal motor neuron as staining in the primary motor cortex also displays a similar pattern of mis-localization (Fig S1A,B). Additionally, cytoplasmic mis-localization was also observed for Importin-ß and TDP-43 (Fig 1B-C,G-H). Interestingly, the loss of nuclear localization of NEMF and TDP-43 was exclusive to neurons in the ventral horn, consistent with lamina IX (Figure S1C,D).

**Figure 1:**
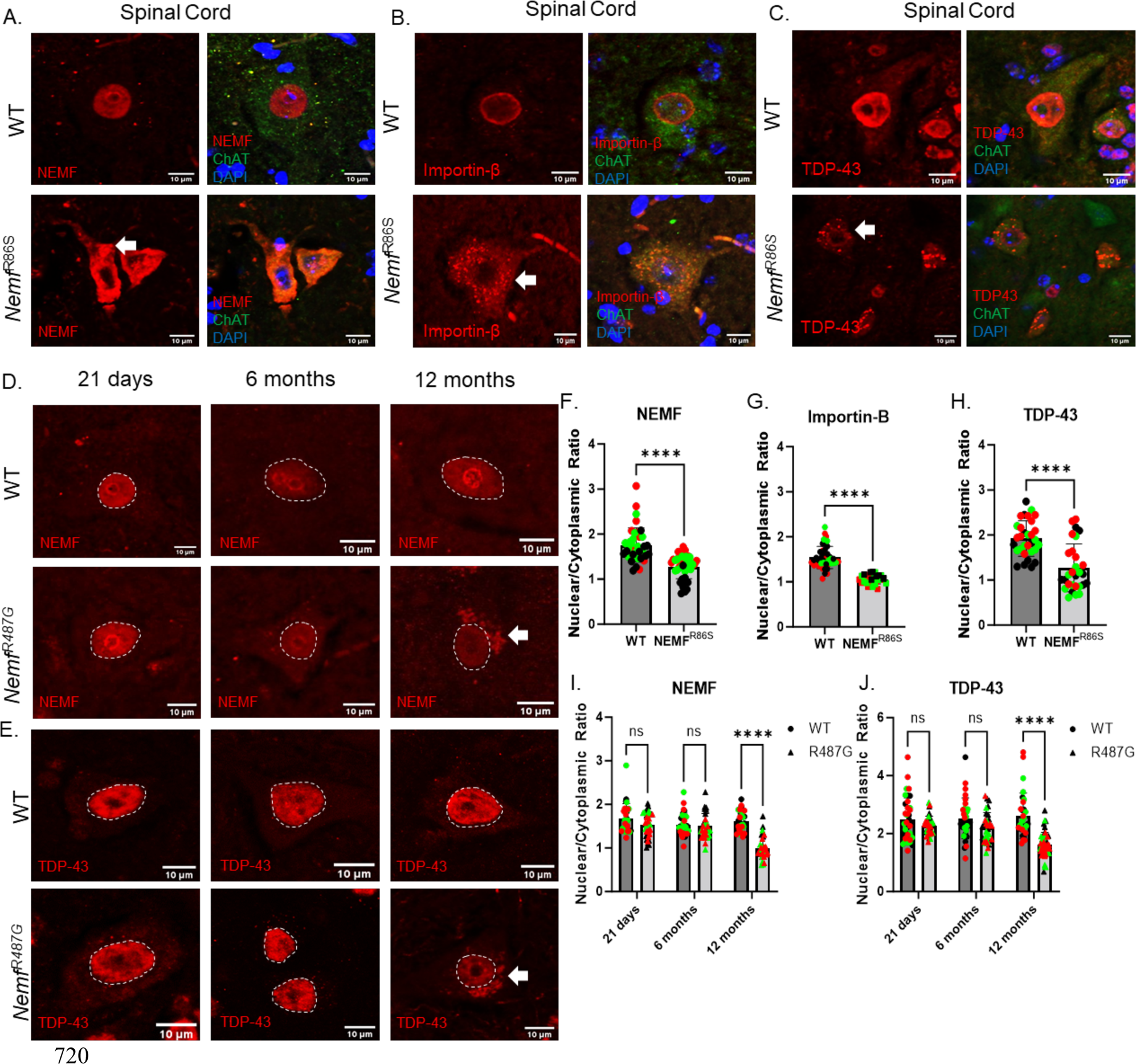
Motor Neuron Degeneration is associated with Progressive Nuclear loss of mutant NEMF in Lumbar Spinal Cord. A-C) Lumbar spinal cords were isolated from 21-day old Wild Type and *Nemf*^R86S^ mice. Motor neurons in the ventral horn were immunostained for the nucleus (DAPI, blue), a motor neuron marker (ChAT, green) and NEMF (A), Importin-β (B), or TDP-43 (C) (red). D-E) Immunostaining of NEMF and TDP-43 (red) in WT, and Nemf^R487G^ lumbar spinal cord motor neurons at 21 days, 6 months and 12 months. F-J) Nuclear/Cytoplasmic ratios of protein of interests in WT and Nemf^R86S^ (F-H) and Nemf^R487G^ (I-J). Data analyzed by unpaired two-tailed t-test (F-H) or two-way ANOVA with Šídák’s multiple comparisons test (I-J). (n=30-44) Arrow indicates cytoplasmic puncta. Scale bars are 10um. (ns p>0.05, **** p<0.0001)

To investigate an expanded temporal scale of the disease progression, we turned to a late onset *Nemf*^R487G^ mouse that also displays a progressive neuromuscular disease with a milder disease phenotype ^20^. The gain of NEMF and TDP-43 cytoplasmic inclusions occurred later in these *Nemf*^R487G^ mice with their appearance beginning at approximately 6 months (Fig 1D). However, the significant nuclear loss of both NEMF and TDP-43 was observed later at 12 months, correlating more closely with disease onset (Fig 1D-E, I-J). Thus, nuclear loss and a toxic cytoplasmic gain of mutant NEMF and TDP-43 occur during disease progression in mice with either of these NEMF mutant alleles.

### *Nemf*^R86S^ exhibits specific Importin-β nuclear import defects

We generated wild type (WT) and *Nemf*^R86S^ mouse embryonic fibroblasts (MEFs) to investigate the mechanisms underlying the NEMF mutant phenotypes. To measure impacts on nuclear import, we used a turboGFP in-frame with a canonical SV-40 Importin-ß nuclear localization signal (GFP-3X-NLS), which we exogenously expressed in *Nemf*^R86S^ MEFs. We found that GFP-3X-NLS appropriately translocated to the nucleus in WT MEFs, but predominantly remained in the cytoplasm in *Nemf*^R86S^ MEFs (Fig 2A,B). A control plasmid with cytoplasmic turboGFP without an NLS showed no difference in localization (Fig 2A,B). We then expressed a Proline-Tyrosine (PY) transportin-1 nuclear localization signal in-frame with mCherry and observed no significant difference in nuclear import (Fig 2C,D). Proper nuclear accumulation of the PY-NLS reporter in both WT and *Nemf*^R86S^ allowed us to create a chimeric protein expressing both a PY-NLS and a pKI nuclear export signal (pKINES) in-frame with an mCherry. Thus, following the initial import into the nucleus, this pKINES-PY-NLS nuclear export reporter would be shuttled out into the cytoplasm and indeed, both the WT and mutant *Nemf*^R86S^ are able to properly export this reporter (Fig 2C,D). Thus, mutant NEMF results in a nuclear transport defect that is specific to the Importin-ß pathway, leaving transportin-1 mediated import and exportin-1 mediated export intact.

**Figure 2:**
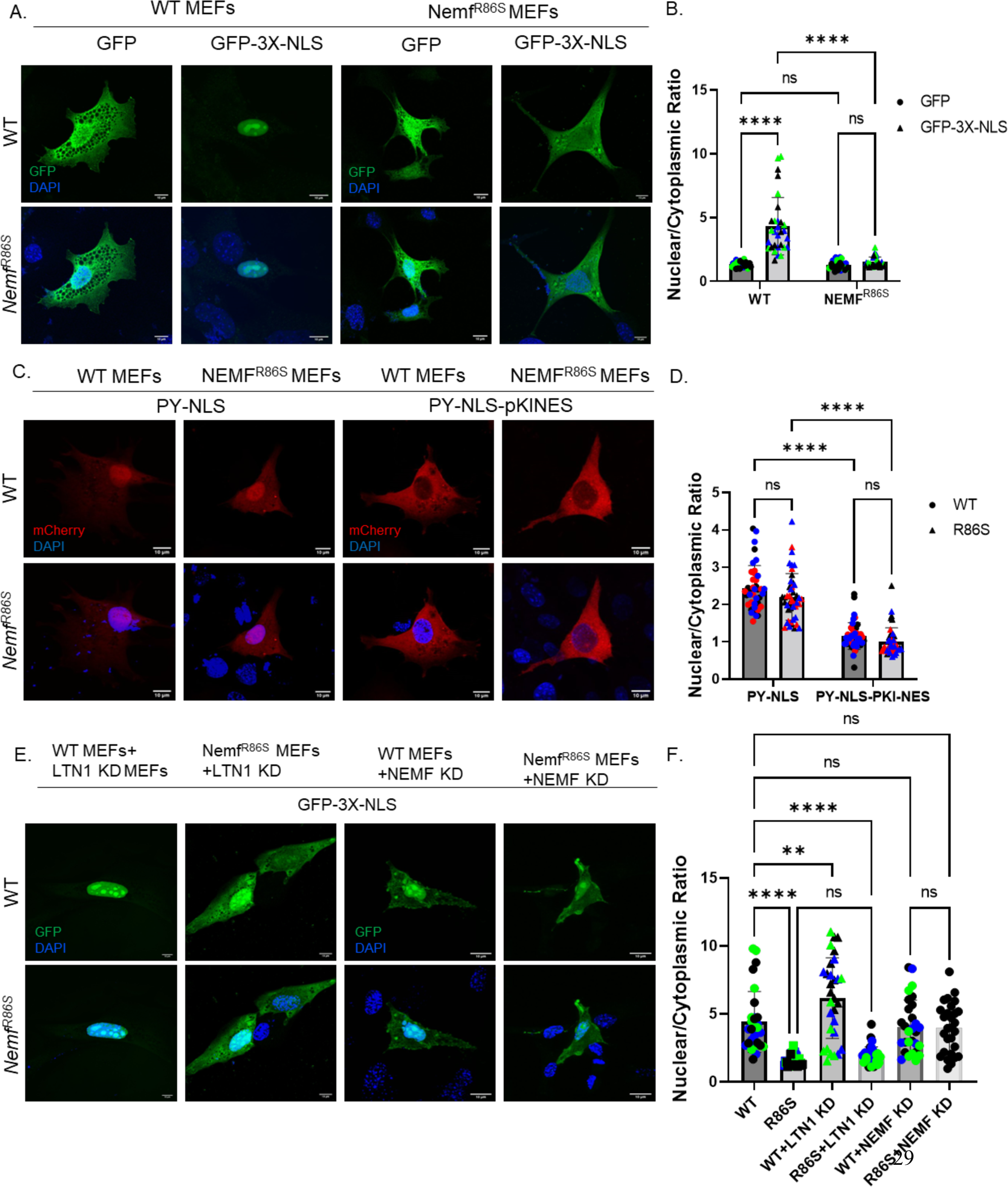
*Nemf*^R86S^ exhibits specific Importin-β nuclear import defects. A) Expression of turbo GFP and turbo GFP in-frame with three Canonical SV-40 Nuclear Localization Signal (3X-NLS) (green) in WT and *Nemf*^R86S^ MEFs. Nuclear stain is DAPI (blue). B) Nuclear/Cytoplasmic Ratios of GFP and GFP-3X-NLS in WT and *Nemf*^R86S^ MEFs. Data Analyzed by two-way ANOVA with Šídák’s multiple comparisons test (n= 30-32) C) Expression of mCherry in frame with a Proline-Tyrosine Transportin-1 Nuclear Localization Signal (PY-NLS) and mCherry in frame with a PY-NLS and a pKI Exportin-1 Nuclear Export Signal (PY-NLS-pKINES) (red) in WT and *Nemf*^R86S^ MEFs. Nuclear stain is DAPI (blue). D) Nuclear/Cytoplasmic Ratios of PY-NLS and PY-NLS-pKINES in WT and *Nemf*^R*86S*^ MEFs. Data analyzed by two-way ANOVA with Šídák’s multiple comparisons test. (n= 40-46) E) Expression of turboGFP-3X-NLS (green) in Ltn1 and NEMF siRNA treated WT and *Nemf*^R86S^ MEFs. Nuclear stain is DAPI (blue) F) Nuclear/Cytoplasmic Ratios of GFP-3X-NLS in each condition. Data analyzed by ordinary one-way ANOVA with Tukey’s multiple comparison test. Scale bars are 10um. (n=28-31, ns p>0.05, **** p<0.0001).

To confirm that this defect is not a result of a passive nuclear import block, we isolated intact nuclei from these MEFs and measured the ability for 20kd, 40kd, and 70kd dextrans to passively diffuse into the nucleus. We observed no significant difference with 20kd and 40kd dextrans but observed a significantly greater nuclear localization of the 70kd dextrans in the *Nemf*^R86S^ MEFs, highlighting a potentially ‘leaky’ nuclear pore in the mutant cells (Fig S2C-E).

To determine whether this Importin-ß defect observed is an indirect effect of RQC dysfunction, we knocked down the E3 ubiquitin ligase *Ltn1* (Fig S2A) and observed there was actually a slight increase in nuclear translocation in WT MEFs, with no difference in the mutant *Nemf*^R86S^ MEFs (Fig 2E,F). Next, we knocked down NEMF (Fig S2B) to determine if this defective nuclear import is either a loss of function or a toxic gain of function phenotype from the mutant form. This *Nemf* knockdown shows that GFP-3X-NLS is capable of translocating into the nucleus, but presents with cytoplasmic puncta (Fig 2E,F).

### *In Situ* Proximity Ligation reveals cytoplasmic NEMF R86S interactions with Importin-ß

Importin-ß has been shown to form cytoplasmic granules under basal conditions as well as under stress ^36^. It has been demonstrated that under conditions of stress such as neurodegenerative disease, both the abundance and size of these granules increases^36^. To identify the cause for this mutant *Nemf*^R86S^ nuclear import defect, we co-stained for Importin-β and NEMF and observed an increase in the size, but not abundance of Importin-β granules (Fig 3A-C). This increase in size was also correlated with an increase in the number of granules colocalizing with NEMF in the R86S cells (Fig 3D). This prompted us to investigate whether the mutant NEMF protein directly interacts with Importin-ß in the cytoplasm. We utilized an in-situ proximity ligation assay using probes for NEMF and Importin-ß and saw both an increase in the mean PLA frequency as well as the mean PLA intensity in the mutant *Nemf*^R86S^ MEFs (Fig 3E-H, S3A). Thus there is an increase in the frequency and strength of interaction between Importin-β and NEMF R86S relative to that with Importin-β and WT NEMF protein. Whether these interactions are a result of direct binding of mutant NEMF protein to Importin-β, or a byproduct of sequestration in a larger complex is yet to be determined.

**Figure 3:**
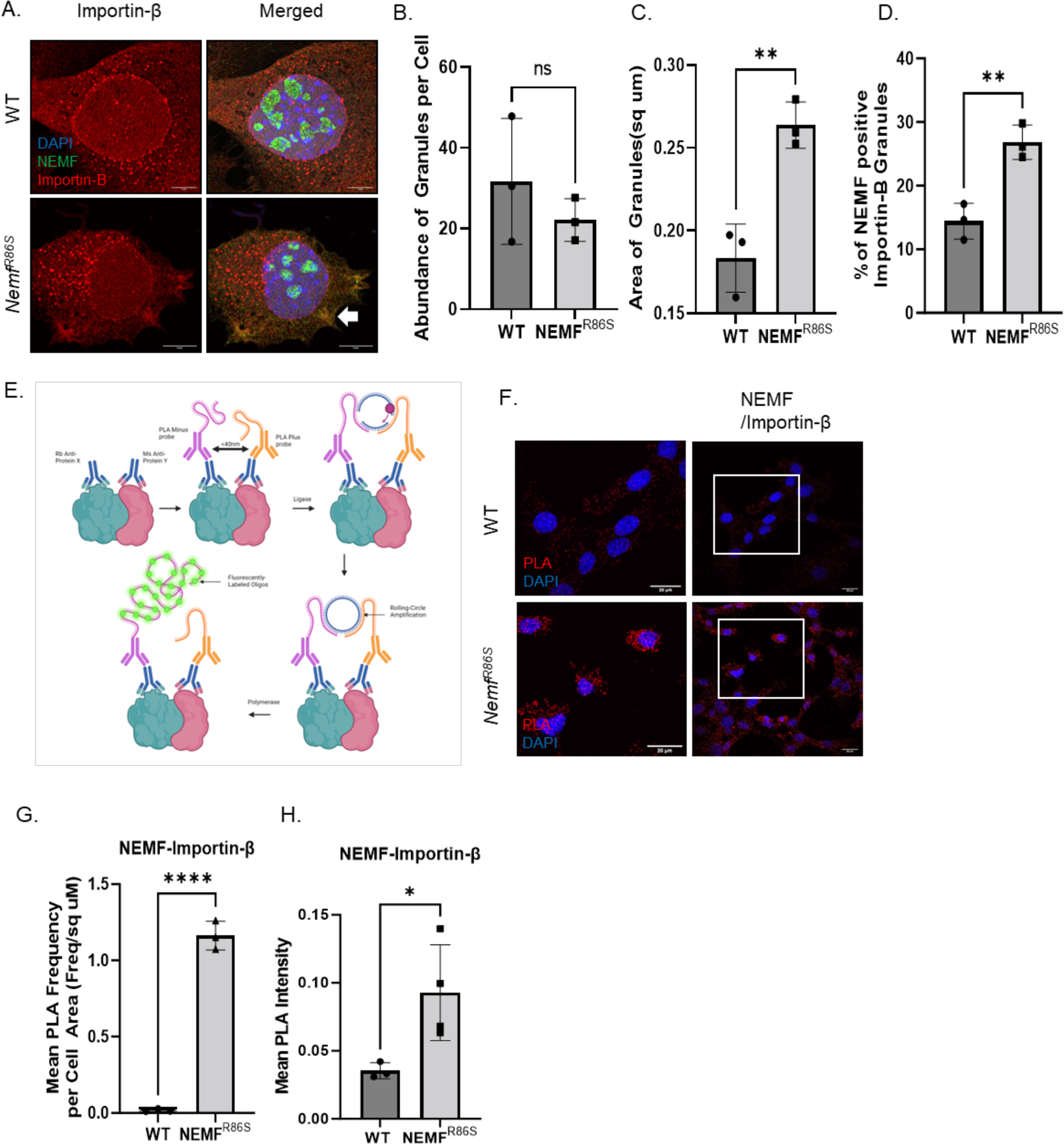
In Situ Proximity Ligation reveals cytoplasmic NEMF R86S interaction with Importin-β. A) Immunofluorescent staining of NEMF (green) and Importin-β (red) in WT and *Nemf*^R86S^ MEFs. Nuclear stain is DAPI (blue). Scale bars are 5um. B-D) Quantification of Importin-β granules (abundance), area of granules (square microns), percent of NEMF positive Importin-β granules (n=3). E) Schematic of PLA adapted from Hegazy et al., 2020 ^66^. F) PLA of NEMF and Importin-B (red). Nuclear stain is DAPI (blue). Scale bars are 20um. G-H) Quantification of Mean PLA frequency and PLA Intensity (n=3). All data was analyzed by unpaired two-tailed t-test. (ns p>0.05, **p<0.01, *p<0.05, ****p<0.0001).

### *Nemf*^R86S^ MEFs display cytoplasmic amyloid-like aggregates

The identification of NCT defects in this mutant *Nemf*^R86S^ model prompted the investigation into the progression of pathology in the form of pathological cytoplasmic aggregates. These *Nemf*^R86S^ MEFs displayed a smaller cell area and a significantly decreased growth rate (Doubling time-WT:17.32h, NEMF R86S: 26.20h) (Fig S4A,B). Immunostaining for NEMF revealed mis-localization of the NEMF R86S protein into cytoplasmic puncta (Fig 4A,B). This phenotype was not uniformly observed, as only a fraction of the cells (20-40%) exhibited this aggregated phenotype (Fig S4D). The mis-localization into cytoplasmic puncta of Importin-ß, TDP-43, Ran, and RanGAP1 suggests that the primary defect of NEMF mutations is a direct dysfunction in nucleocytoplasmic transport (Fig 4A, C-F). Additionally, various nucleoporins were also observed to be sequestered within these cytoplasmic puncta (Nup50, Nup98, Nup153) (Fig S4C). AmyloGlo (Biosensis) labeling also revealed a significant increase in the amyloid-like signal in the cytoplasm (Fig S4E,F) in the mutant MEFs compared to the WT MEFs. As TDP-43 was found to be mis-localized, pathological TDP-43 was identified through phospho-TDP-43 immunostaining (Fig 4G) and was found to be greatly increased in mutant cells. We then biochemically isolated TDP-43 using a sarkosyl insoluble fractionation ^37^ in WT and *Nemf*^R86S^ MEFs and we found an increase in levels of insoluble TDP-43 in mutant cells (Fig 4H,I, S4E). Immunostaining of spinal motor neurons in WT and mutant *Nemf*^R86S^ mice revealed cytoplasmic phospho-TDP-43 aggregation exclusive to these ChAT-positive motor neurons (Fig 4J). Subsequent staining of the late onset mutant *Nemf*^R487G^ spinal motor neurons revealed some phospho-TDP-43 positive motor neurons at 6 months, with most motor neurons positive at 12 months (Fig S5).

**Figure 4:**
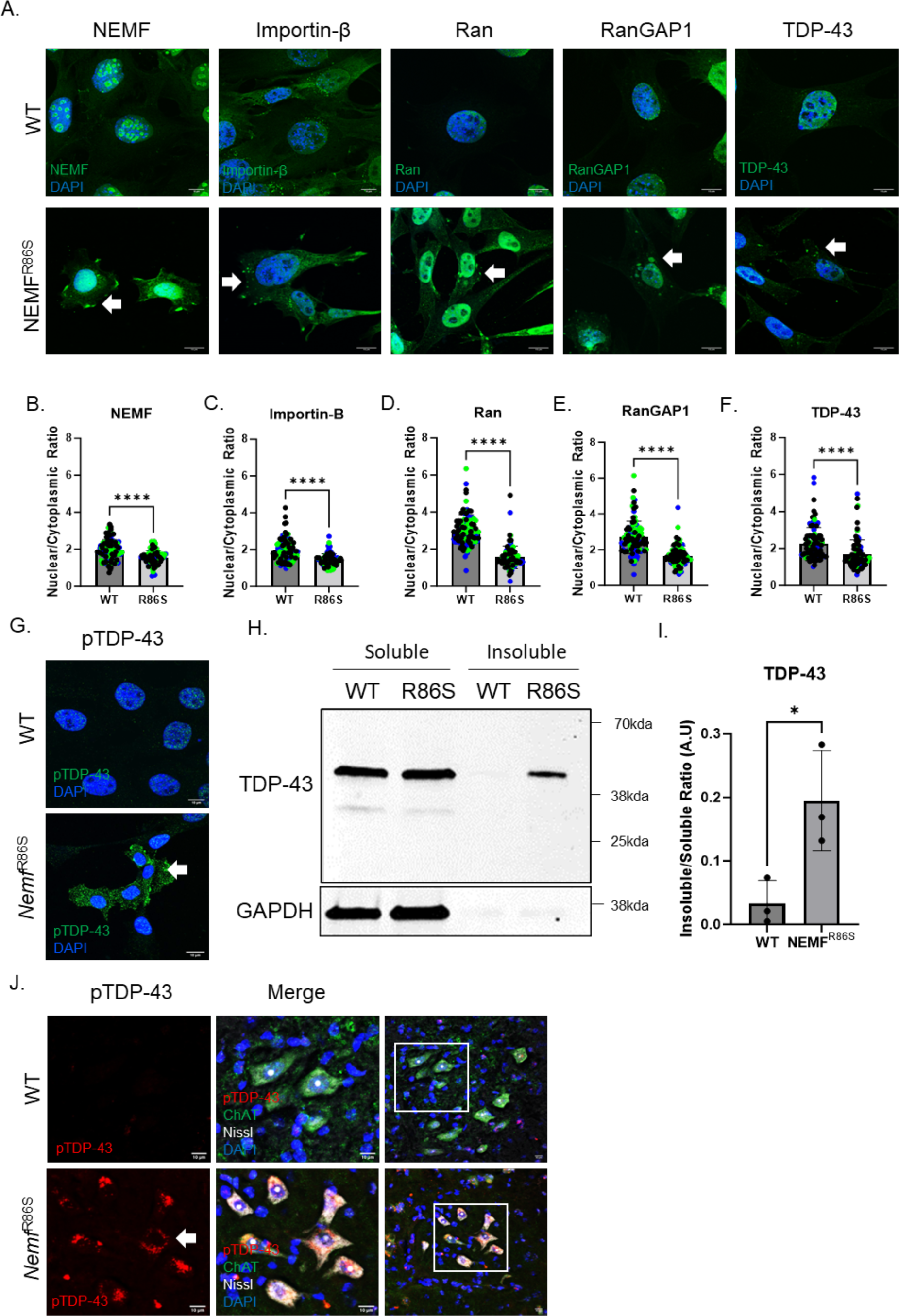
*Nemf*^R86S^ MEFs display cytoplasmic amyloid-like aggregates. A) Immunofluorescent Staining of NEMF, Importin-β, RanGAP1, Ran, and TDP-43 (green) in WT and *Nemf*^R86S^ MEFs. Nuclei Labeled with DAPI (blue). B-F) Quantification of Nuclear/Cytoplasmic Ratio of indicated proteins (n=100). Data analyzed by unpaired two-tailed t-test. G) Immunofluorescent Staining of pTDP-43 in WT and *Nemf*^R86S^ MEFs. H) Western Blot Analysis of sarkosyl soluble and insoluble TDP-43 in WT and *Nemf*^R86S^ MEFs I) Ratio of Soluble versus Insoluble TDP-43 protein levels (n=3). J) Immunofluorescent staining of phospho-TDP-43 (red), ChAT (green), Nissl (white) in WT, and *Nemf*^R487G^ lumbar spinal cord motor neurons at 21 days. Arrow indicates cytoplasmic puncta. Scale bars are 10um. (*p<0.05, ****p<0.0001).

Taken together, these findings demonstrate that NEMF mutations cause cytoplasmic protein aggregates, leading to the sequestration of nuclear transport factors and TDP-43 pathology.

### *Nemf*^R86S^ mice displays altered expression of ALS-linked genes

The nuclear loss of TDP-43 prompted us to examine transcriptional profiles in this mutant model to determine whether there were similar transcriptional changes as those seen in ALS/FTD with TDP-43 dysfunction. Indeed, a subset of ALS- and AD-linked genes displayed altered expression in the mutant MEFs, with *Stmn2* and *Apoe* being highly down- and upregulated, respectively (Fig 5A,B). qPCR validation in MEFs shows that both transcripts are significantly dysregulated in mutant, with significant dysregulation also found in lumbar spinal cord tissue, but not in brain tissue (Fig 5C-D). Other notable genes found to be dysregulated include *Fig4*, *Bax,* and *Sorl1*, which were significantly dysregulated *in vitro* but not *in vivo* (Fig 5F, S6E-F). However, this discrepancy could be attributed to the non-specificity of bulk tissue RNA extraction. Another gene commonly dysregulated in neurodegeneration, *Chmp2b,* was downregulated in all tissue types, but was not significant (Fig 5E). To validate RNAseq data to qPCR data, simple linear regressions were used to model fold change in RNA-seq data and qPCR data, showing significant correlations in MEFs and spinal cord tissue, but not brain tissue (Fig S7C). An additional upregulated long noncoding RNA, *Meg3*, has been implicated in necroptosis in Alzheimer’s disease (Fig 5B)^38^. Gene ontology and pathway analysis of the RNA-seq dataset of WT and mutant *Nemf*^R86S^ MEFs revealed biological processes involved in cell adhesion, skeletal system development, and negative regulation of cell proliferation (Fig S6). An investigation into STMN2 protein levels revealed STMN2 protein loss *in vitro* and *in vivo*, and specifically in the spinal cord, but not the brain, consistent with RNA downregulation (Fig 5G-L). Collectively, these observations, both *in vitro* and *in vivo,* indicate that a primary pathology triggered by canonical nuclear import defects results in TDP-43 pathology and downstream transcriptional dysregulation in sporadic forms of ALS-FTD and neurodegenerative disease.

**Figure 5:**
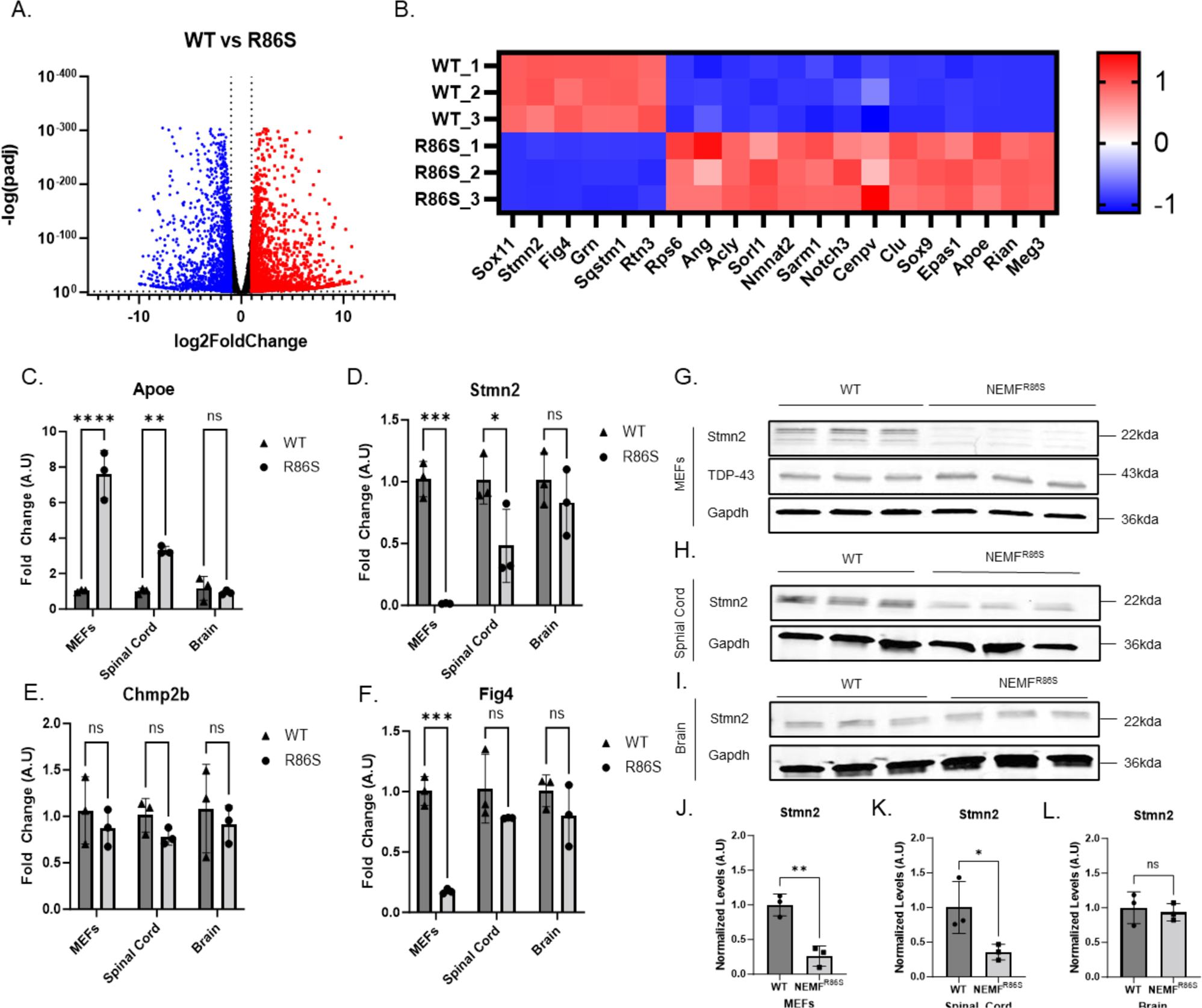
*Nemf*^R86S^ mice display altered expression of ALS-linked genes. A) Volcano plot of upregulated (red) and downregulated (blue) genes. (Log2foldchange threshold <-1, >1, p<0.05). B) Heatmap of significantly dysregulated genes commonly observed in neurodegeneration. (C-F) qPCR relative fold change of *Stmn2, Apoe, Chmp2b, and Fig4* in MEFs, spinal cord, brain in WT and *Nemf*^R86S^ mice. Data analyzed by two-way ANOVA with Šídák’s multiple comparisons test. (n=3). G) Western Blot Analysis of STMN2 protein levels in MEFs, Spinal Cord, and Brain. J-L) Quantification of STMN2 protein levels normalized to GAPDH levels. Data analyzed by unpaired two-tailed t-test (n=3) (ns p>0.05, *p<0.05, **, p<0.01, *** p<0.001, **** p<0.0001)

### A transient nuclear import block recapitulates transcriptional downregulation of *Stmn2*

To investigate if the Importin-β pathway itself recapitulates the phenotypes seen in the NEMF mutant, we targeted the nuclear import pathway through a small molecular inhibitor of Importin-β, importazole (IPZ) ^39^. We found that transiently blocking Importin-β nuclear import results in the mis-localization of TDP-43 with the presence of cytoplasmic phospho-TDP-43 (Fig 6A,B) and in the downregulation of *Stmn2* transcript, with no alteration of *Apoe* transcripts (Fig 6C,D). To confirm that *Stmn2* downregulation was in fact downstream of blocking nuclear import, and not simply a side effect of importazole treatment, we also treated MEFs with ivermectin (IVM), a known Importin-β antagonist, and observed a similar downregulation of *Stmn2* transcript, with no *Apoe* dysregulation (Fig 6C,D). A cell viability assay was used to determine sub-toxic levels of importazole (20uM) and ivermectin (10uM) following a 48h treatment (Fig S6A,B). To further ensure that the downregulation observed was not a result of a decrease in RNA quality, as has previously been described to be a confounding factor ^40^, RNA quality was checked and confirmed to be of the highest quality, with RINe values ranging from 9.4-9.7 across all samples (Fig S7). Following these observations, Importazole-treated RNA samples were sent for bulk RNA sequencing to determine which transcripts were similarly dysregulated to our *Nemf*^R86S^ MEFs (Fig 6E). We found 20 commonly dysregulated genes in both conditions, with 15 of those genes similarly downregulated, and *Stmn2* being the only ALS-linked gene commonly downregulated (Fig 6F,G). A gene ontology analysis also described cell adhesion as being the most significant dysregulated pathway (similar to the *Nemf*^R86S^ MEFs) with collagen fibril organization, extracellular matrix organization, and many cholesterol and lipid processes also observed as dysregulated pathways (Fig S6C,D).

**Figure 6:**
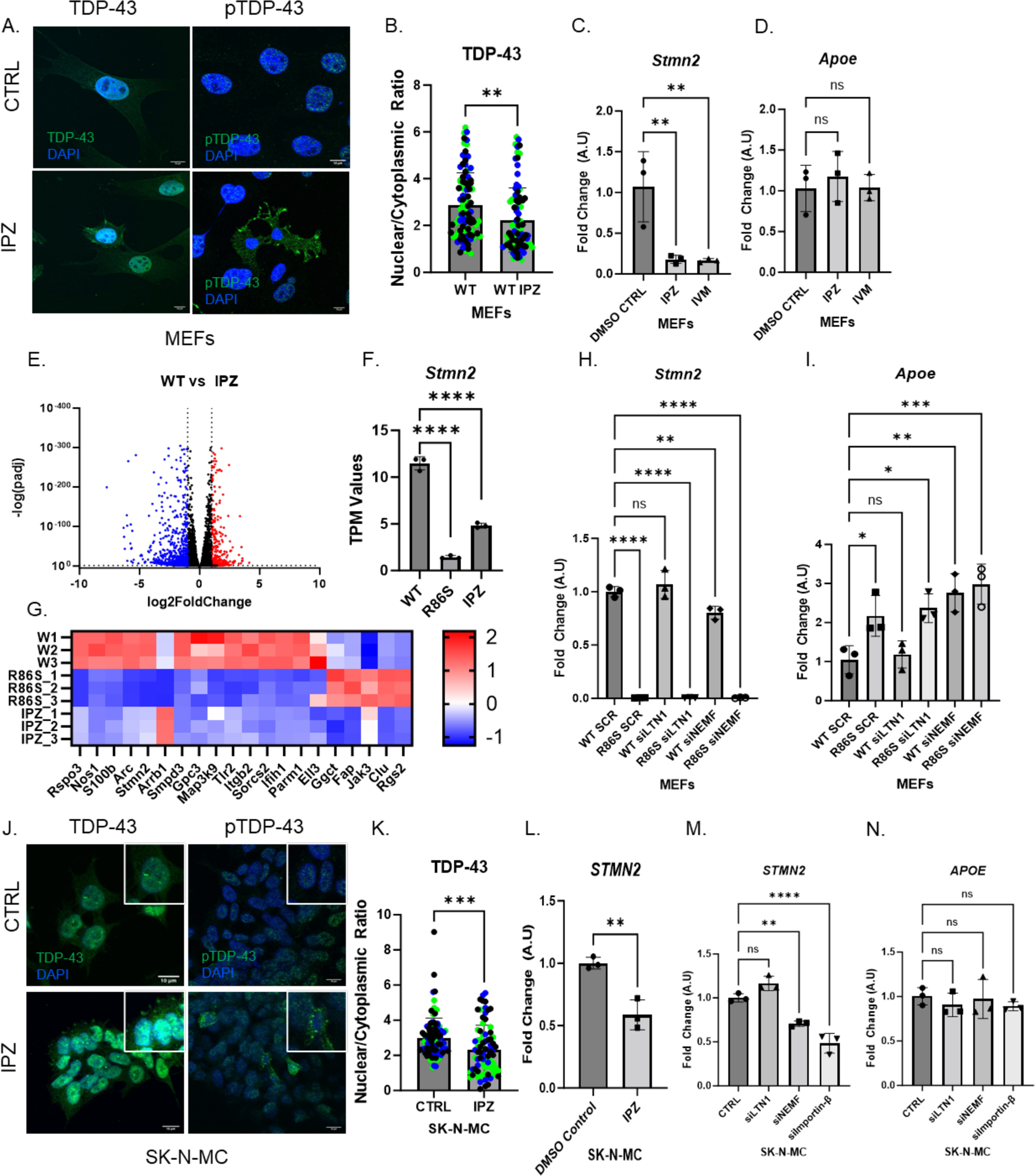
A transient nuclear import block recapitulates transcriptional downregulation of *Stmn2*. A) Immunofluorescent staining of TDP-43 and pTDP-43 in WT and Importazole-treated (IPZ) MEFs. B) Quantification of Nuclear/Cytoplasmic Ratio of TDP-43 (n=92-99). Data analyzed by unpaired two-tailed t-test. C-D) qPCR relative fold change in DMSO Control and IPZ and Ivermectin (IVM) treated MEFs for indicated genes (n=3). E) Volcano plot of upregulated (red) and downregulated (blue) genes in IPZ-treated MEFs bulk RNA. (Log2foldchange threshold <-1, >1, p<0.05). F) *Stmn2* transcripts per million values for WT, *Nemf*^R86S^, and IPZ-treated MEFs. G) Heatmap of significant commonly dysregulated genes in WT, *Nemf*^R86S^, and IPZ-treated MEFs. (n=3) H-I) qPCR relative fold change of *Stmn2* or *APOE* in WT or *Nemf*^R86S^ MEFs treated with *Ltn1* or *Nemf* siRNAs (n=3). J) Immunofluorescent staining of TDP-43 and pTDP-43 in WT and Importazole-treated (IPZ) SK-N-MC (neuroblastoma) cells. K) Quantification of Nuclear/Cytoplasmic Ratio of TDP-43 (n=80-96). L) qPCR relative fold change in DMSO Control and Importazole (IPZ) treated SK-N-MC cells for *STMN2* (n=3). M-N) qPCR relative fold change of *STMN2* or *APOE* treated with *LTN1*, *NEMF*, or *Importin-β* siRNAs in SK-N-MC cells (n=3). Data analyzed by one-way ANOVA with Tukey’s multiple comparison test. (ns p>0.05, *p<0.05, **, p<0.01, *** p<0.001, **** p<0.0001)

Using *Apoe* and *Stmn2* dysregulation as a marker for transcriptional dysregulation, we targeted *Nemf* and *Ltn1* by siRNA knockdown (Fig S8A,B) and saw that *Nemf* knockdown causes a significant reduction in *Stmn2* transcript and an almost 3-fold increase in *Apoe* transcript, comparable to the mutant mouse model (Fig 6H,I). In comparison, *Ltn1* knockdown samples did not perturb *Stmn2* and *Apoe* transcriptional regulation (Fig 6H,I).

To confirm that this transcriptional dysregulation can be recapitulated in human models, we treated SK-N-MC neuroblastoma cells with importazole for 48h and observed an increase in the cytoplasmic mis-localization of TDP-43 with the presence of cytoplasmic phospho-TDP-43 (Fig 6J,K). This TDP-43 mis-localization was accompanied by a ∼50% reduction in *STMN2* transcripts (Fig 6L). We then proceeded to knock down *NEMF* (SDCCAG1) and *LTN1* (RNF160) in HEK293 and SK-N-MC cells (Fig S8A,B) and observed a similar dysregulation of *STMN2* but not *APOE* transcripts (Fig 6M,N, Fig S8C,D). We found a downregulation of full-length *STMN2* transcripts in si*NEMF* treated HEK293 and SK-N-MC cells, but not si*LTN1* treated samples (Fig 6M, Fig S8C). We also observed an increase in *APOE* transcripts in si*NEMF* treated HEK293 cells (Fig S8D), but not in si*LTN1* treated HEK293 cells or in both treatments in SK-N-MC cells (Fig 6N, Fig S8D). To further elucidate how nuclear import mechanisms can dysregulate *STMN2* transcriptional regulation, we targeted nuclear import pathways through siRNA knockdown (*Importin-β* (KPNB), *Importin-α* (*KPNA*), and *Transportin-1 (TPNO1*)) in HEK293 cells (Fig S8A) and *Importin-β* in SK-N-MC cells (Fig S8B). Interestingly, we see a downregulation of *STMN2* transcripts when Importin-β nuclear import is inhibited either by *Importin-β* siRNA-treated or IPZ-treated samples in both HEK293 and SK-N-MC cells (Fig 6L,M, S8C,E). Furthermore, only knockdown of *Importin-α* in HEK293 resulted in a significant upregulation of APOE (Fig 6D). Altogether, our results show that targeting nuclear import pathways, as well as *Nemf,* can recapitulate *Stmn2* downregulation in both mice and human cell lines.

## Discussion

A common pathological hallmark of neurodegeneration is the accumulation of cytoplasmic amyloid-like aggregated proteins ^16,41,42^. Our data, utilizing two independent single nucleotide polymorphisms (R86S and R487G) in *Nemf*, demonstrate that disrupted Importin-βnuclear import results in the mis-localization of nuclear proteins to the cytoplasm. Importantly, this disruption of NCT is specific to Importin-β nuclear import.

Recent evidence indicates that cells actively promote the sequestration of misfolded proteins into aggregates, which are further compartmentalized within distinct cellular compartments ^43–45^. NIRs have been shown to operate in the cytoplasm and disaggregate NLS-bearing cargoes to prevent deleterious phase transitions ^46^. This ability to inhibit and reverse both physiological and deleterious phase transitions has been implicated in multiple forms of neurodegenerative diseases, wherein RNA-binding proteins or Prion-like domains are lost from the nucleus and accumulate in the cytoplasm ^46–54^. This is demonstrated in the role of Importin-β cooperating with Importin-α to prevent and reverse TDP-43 condensation and fibrillization ^46,48^. The potential of these NIRs to serve as cytoplasmic chaperones and restore the nuclear function of RNA-binding proteins highlights their therapeutic potential ^55^.

It is difficult to establish whether the mutant NEMF R86S is pathologically interacting with Importin-β directly or is indirectly associating with Importin-β through aggregate-induced sequestration. However, the absence of nuclear import defects in an aggregate prone *Ltn1* knockdown model highlights a specificity for NEMF in these observed import defects. This fact, along with the greater association of the NEMF R86S protein with Importin-β through PLA, highlights a role for NEMF interactions with Importin-β. The early onset of disease in the *Nemf*^R86S^ mutant mice suggests that the cytoplasmic accumulation of NEMF and TDP-43 may be a result of a greater propensity to mis-localize in the R86S variant when compared to the R487G, whether through direct or indirect mechanisms. Importantly, the mis-localization of NEMF, Importin-β, Ran, RanGAP1, and TDP-43 in the mutant *Nemf*^R86S^ MEFs is observed broadly throughout the cell population whereas the presence of cytoplasmic puncta is only observed in a subset of the cell population. This indicates that the nuclear import defects may precede the accumulation and sequestration of NTRs in the cytoplasm. Furthermore, the presence of insoluble TDP-43 and phospho-TDP-43 in the *Nemf*^R86S^ MEFs and spinal cord motor neurons is consistent with TDP-43 pathology in ALS and FTD/FTLD ^56,57^.

The loss of *Stmn2* transcript in our *Nemf*^R86S^ mouse model suggests either that *Stmn2* is downregulated directly because of defects in nuclear import, that its down regulation is a result of the mis-localization of TDP-43 itself, or some combination of the two. Our observations of pathological TDP-43 insoluble aggregates and a downregulation of *Stmn2* mRNA and protein after a nuclear import block indicates a direct causal role of defective nuclear import in both of these pathologies. Reduced *Stmn2* expression has been implicated in human TDP-43 pathologies in ALS and AD, as well as in Parkinson’s disease patients and FTLD-tau patients ^40,58–61^. Furthermore, the genetic loss of *Stmn2* alone in mice results in motor neuropathy ^62,63^. Additional dysregulation of transcripts, such as the upregulation of *Meg3* and *Apoe*, as well as the downregulation of *Fig4*, are seen in our model and mirror changes seen in neurodegenerative disease^64,65^. Our investigations show that by disrupting *NEMF* and *Importin-β* pathways through knockdown in human cell lines, we can induce a significant decrease in *Stmn2* transcript. Interestingly, knockdown of *NEMF* and *Importin-α*, but not *Importin-β* causes an upregulation of *APOE* transcripts, again mimicking similar phenotypes observed in mice.

We have demonstrated that there is a direct impact of Importin-β nuclear import dysregulation on multiple phenotypes seen in neurodegeneration through the following: (1) NEMF-specific dysregulation of Importin-β nuclear import (but not of Transportin-1 import), (2) mis-localization and aggregation of NEMF, TDP43, Ran, RanGAP1 and Importin-β, (3) amyloid-like aggregation, (4) PLA cytoplasmic association between NEMF R86S and Importin-β, (5) cytoplasmic accumulation of NEMF, Importin-β, and TDP43 in *Nemf* mutant mouse spinal cord, (6) alterations in ALS and AD-related transcripts including *Stmn2*, *Apoe*, *Fig4*, and *Sorl1*, and (7) transient Importin-β-specific nuclear import blockage or *Nemf* siRNA knockdown recapitulate these transcript alterations. These dysfunctions are not, however, recapitulated upon knockdown of NEMF’s RQC partner LTN1, indicating a specific, non-RQC role for NEMF in impairing Importin-β nuclear import. Collectively, our data suggests that Importin-β nuclear import dysfunction serves as a primary upstream pathomechanism for TDP-43-mislocalization and phosphorylation, as well as transcriptional dysfunction seen in post-mortem ALS patient brain and spinal cords.

## Acknowledgments

We wish to thank current and former members of the Sher lab for critical discussions and advice. We wish to thank Dr. Gregory Cox (Jackson Laboratory and University of Maine) for help with providing NEMF mutant mice and with suggestions for research strategies. We thank Dr. Joshua Dubnau for experimental design advice. We thank Dr. Leonard Petrucelli (Mayo Clinic) for the kind gift of the mouse-reactive phospho-TDP43 antibody. We thank Wendy Ackmentin for her incredible help on all aspects of keeping the lab facilities running.

## Experimental Procedures

### WT-NEMF and R86S-NEMF Mouse Embryonic Fibroblasts (MEFs) Culture

WT NEMF and R86S-NEMF MEFs were maintained in Dulbecco’s Modified Eagle’s Medium (10% FBS, Glutamine 1X Glutamax), PenStrep) at 37degC, 5% CO2 and passaged every two days with Trypsin-EDTA (0.05%). Reagents used indicated in Supplementary Table.

### HEK293 Culture

HEK293 cells were maintained in Dulbecco’s Modified Eagle’s Medium (10% FBS, 1mM Glutamax, PenStrep, Non-essential amino acids) at 37degC, 5% CO2 and passaged every two-four days with Trypsin-EDTA (0.05%). Reagents used indicated in Supplementary Table.

### SK-N-MC Culture

SK-N-MC cells (ATCC) were cultured according to the vendor’s protocol. In brief, cells were maintained in ATCC-formulated Eagle’s Minimum Essential Medium (10% FBS, PenStrep) at 37C in 5% CO2. Reagents used indicated in Supplementary Table.

### Microplate MEF Plating

Coverslips (Carolina Biological Supply) were washed with 70% ethanol in a sterile petri dish to sterilize and then allowed to air dry for 30mins. Using a 24 well microplate, the sterilized coverslips were then placed inside each well, and then 1mL of the 0.1%(w/v) gelatin (Sigma) was added and incubated at 37degC for 1hr. The excess gelatin was aspirated with a vacuum and the coverslips were again incubated at 37degC for 1hr.

### Plasmid and SiRNA Transfections

Transfection of plasmids and siRNAs (Supplementary Table) were performed using Lipofectamine 3000 and Lipofectamine 2000 (Life Technologies) following manufacturer’s instructions. Experiments were performed 48h after transfections of plasmids and siRNAs.

### Nuclear Import Block

Cells were treated with 20uM and 10uM of importazole (Sigma) and ivermectin (Sigma) respectively for 48 hours at 37degC 5% CO2.

### Immunofluorescent Staining

Cells were chemically fixed in 4% paraformaldehyde in PBS(1X). Each well was then rinsed with PBS(1X). The cells were then permeabilized with 0.1% TritonX-100/PBS for 10mins at room temperature. The cells were then blocked at room temperature with 5% NGS/PBS/0.1% Tween-20. The cells were then incubated with the respective primary antibody in 5% NGS/0.1% Tween-20/PBS overnight at 4degC. The primary antibodies used are listed in Supplementary Table. The next day, the cells were rinsed 3 times for 5mins with 0.1% Tween-20/PBS(1X). The cells are then incubated with the respective secondary antibodies in 5% NGS/0.1% Tween-20/PBS for 2hrs in the dark at room temperature. The secondary antibodies used are listed in Supplementary Table. The cells were then rinsed 3 times for 5mins with 0.1% Tween-20/PBS and stored temporarily with 200µL of PBS(1X) in the dark at 4degC. Using sterile slides, 20-25µL of Prolong glass with NucBlue (Thermo Fisher) was added to the slide. The liquid was then aspirated from the well and using an SE 5 tissue curved forcep, the coverslips were gently picked up and then placed with the cell layer (top) down on the slide. The slide is then cured in the dark at 4degC for 24hrs. Confocal images were taken by an Olympus FV1000 Laser Scanning Confocal.

### Growth Assay

Cells were seeded on a 24 well plate at a cell density of 10,000 cells per well. Using Agilent Lionheart Imager, three regions of interests in the well were imaged for 48 hours every 15mins. Cell growth was standardized by starting cell number.

### RIPA Lysis Protein Extractions

Cells were washed twice with 5mL of pre-chilled PBS(1X). 500µL of RIPA lysis buffer (Sigma) supplemented with protease inhibitor was added to the flask and then incubated on ice for 5mins. The cells were scraped and then transferred to a 1.5mL microtube. The lysate was then agitated for 30mins at 4deg C and then centrifuged for 20mins at 12,000rpms at 4deg C. The supernatant was then transferred to a pre-chilled microtube and stored at −80deg C.

### Sarkosyl Insoluble Fractionation

Pathological TDP-43 aggregates were biochemically isolated as previously described in Manuela et al 2019 ^37^. In brief, cells grown in a 6 well plate were washed twice in 1ml of PBS (1X). 100ul of 1X HS Buffer (10mM Tris, pH 7.4, 15mM NaCl, 0.5mM EDTA, 1mM DTT, protease and phosphatase inhibitors) supplemented with 0.5% Sarkosyl (Sigma) and Benzonase Endonuclease (12mM MgCl2 with 250U/Sample Benzonase) (Sigma) was added to the wells and the cells were scraped into a 1.5ml microtube. The wells were washed 1X with 1X HS Buffer, 0.5% Sarkosyl and added to the microtube. 200ul of 2X HS Buffer, 4% Sarkossyl was added to the lysate and incubated on ice for 45mins with vortexing in 10min intervals. Following cell lysis, 200ul of ice cold 1X HS Buffer, 0.5% Sarkosyl was added the sample and then centrifuged at max speed for 20mins at RT. The supernatant was collected as the soluble fraction, and the pellet was washed twice with PBS 1X. Pellets were frozen at −80degC until SDS PAGE.

### Pierce BCA Protein Assay

The standards were prepared according to the Test Tube protocol provided by the Pierce BCA (Thermo Fisher). 10µL of each standard was pipetted into appropriately labeled 500µL microtubes and repeated for each label. 200µL of working reagent was added to each tube and incubated in a water bath at 37degC for 30mins. The standards were analyzed on the NanoDrop at 562nm to establish a standard curve and the samples were then analyzed on the curve.

### SDS-PAGE

10µg of lysate was prepared with 50mM dTT (Biorad) and 4X laemli buffer (Biorad). The solution was vortexed quickly and boiled for 5mins at 95degC. The lysates were then chilled briefly on ice and then loaded onto a Stain-Free mini-protean 10 well pre-cast gel (BioRad) and mini-protean tank (BioRad) with a Chameleon 800 MW ladder (Licor). The gels were run at 200V for approximately 45mins in Tris/Glycine/SDS Running Buffer (BioRad). Gels were then equilibrated in transfer buffer (20% methanol) (BioRad) and total protein was imaged using a ChemiDoc Imaging System (BioRad).

### Semi-dry Transfer (PVDF .2um)

The PVDF membrane was pre-wet in methanol, then rinsed with ultra-pure water. The membrane and 6 filter papers are equilibrated for 20mins in transfer buffer. A gel sandwich is then prepared with 3 filter papers, gel, membrane, 3 filter papers, respectively. The transfer is then run for 90mins, with the current monitored to maintain 80mA-240mA. After the transfer, the membrane is removed and allowed to dry completely. The membrane is then re-activated with methanol, and rinsed once with TBS (1X), then stored in TBS(1X) in the fridge until immunoblotting.

### Immunoblotting

The membrane is rinsed with TBS(1X) and then incubated with TBS-Based Odyssey blocking buffer for 1hr at room temperature with gentle rocking. The blocking buffer is then discarded, and the membrane is incubated in blocking buffer supplemented with 0.1% Tween-20 and primary antibody overnight with gentle rocking. The primary antibodies used are listed in the Supplementary Table. The membrane is then washed with TBS-T (0.1% Tween-20) 3 times for 5mins with gentle shaking. The membrane is then incubated with blocking buffer (0.1% Tween-20) and the respective secondary antibody at room temperature for 2hrs with gentle rocking. The secondary antibodies used are listed in the Supplementary Table. The membrane is then washed with TBS-T 3 times for 5mins. The membrane is then rinsed with TBS(1X) and imaged with the Odyssey Scanner (Licor).

### Mouse strains, husbandry, and genotyping

All mouse husbandry and procedures were reviewed and approved by the Institutional Animal Care and Use Committee at Stony Brook University and were carried out according to the NIH Guide for Care and Use of Laboratory Animals. Tail tissue was lysed in proteinase K at 55deg C overnight and extracted DNA was used for genotyping. Genotyping for B6J-*Nemf*^R86S^ was performed via PCR using the following primers: forward primer specific to wild-type allele: 5′-AACATTTGAAGAGTCGGGGA-3′; forward primer specific to mutant allele: 5′-AACATTTGAAGAGTCGGGGT-3′; reverse primer common for both alleles: 5′-GCAGGTGGATGGTAGCAACG-3′. Similarly, for the genotyping of the B6J-*Nemf* ^R487G^ mice the following primers were used: forward primer specific to wild-type allele: 5′-TGCTGCTAAAAAAACCCGGA-3′; forward primer specific to mutant allele: 5′-TGCTGCTAAAAAAACCCGGG-3′; reverse primer common for both alleles: 5′-AAAGCCCTTGCTGCAAAGCC-3′.

### Spinal Cord Isolation

Mice were euthanized by carbon dioxide inhalation and immediately exsanguinated by cardiac puncture using a 1mL syringe with a 25g needle. Using small surgical scissors and forceps, the dorsal skin was removed. Next, the location of the atlanto-occipital joint was visualized by repeated flexion and extension of the spinal column and palpation. Using large scissors, animals were decapitated at the atlanto-occipital joint, exposing the cervical spinal cord. Following decapitation, a clean transverse cut was made through the lumbar portion of the spine, cranial to the iliac crest. Moving from caudal to cranial, the vertebral column was dissected from the ventral portion. The entire vertebral column was rinsed with PBS 1X and placed in a 15mL conical tube with 4% PFA for 20mins at 4degC. During this stage, skull cap is removed with brain gently teased from the cranial cavity in a rostro caudal direction. Micro-dissection forceps are used to gently tease apart the cranial nerves. The brain is then rinsed in PBS 1X and placed in a 15ml conical tube with 4% PFA at 4degC with agitation overnight. The vertebral column stored in 4% PFA is then placed into a 100mm petri dish with PBS 1X. Moving from caudal to cranial, fine-tipped offset bone nippers were used to reveal the dorsal aspect of the spinal cord. The vertebral column is then held vertically, and the spinal cord is eased from the spinal canal using micro-dissection forceps and placed into the 100mm petri dish. The spinal cord was then placed into a new 15ml conical tube with 4% PFA at 4degC with agitation overnight. The following day, the brain and spinal cord are both transferred to 30% sucrose (w/v) for an additional 24h. The following day, the spinal cord is sectioned above the lumbar enlargement region with microscissors. The brain, the lumbar region and the thoracic and cervical regions are both placed in optimal cutting temperature (OCT) and stored in −80degC.

### Cryosectioning

Lumbar spinal cords and cervical and thoracic spinal cords stored in OCT were placed in −20degC for one hour prior to cryosectioning to allow for the temperature to equilibrate. Spinal cords were then sectioned transversely in the cranial to caudal direction at 30um slices and placed free-floating in a 24 well microplate filled with PBS 1X at 20 sections per well. Sections were allowed to equilibrate in PBS 1X for 1 hour. Sections were then rinsed with PBS 1X to remove residual OCT and were then redistributed to a 48 well plate in PBS for immunoblotting.

### Spinal Cord Immunoblotting

Spinal Cords were rinsed with PBS (1X). Spinal Cords were then permeabilized in 0.3% TritonX-100/PBS 1 hour at room temperature with agitation. The spinal cords were then blocked at room temperature with 5% NGS/PBS/0.1% TritonX-100. The spinal cords were then incubated with the respective primary antibody in 5% NGS/PBS/0.1% TritonX-100 overnight at 4degC. The primary antibodies used are listed in Supplementary Table. The next day, the spinal cords were washed 3 times for 15mins with 0.1% TritonX-100/PBS(1X). The spinal cords are then incubated with the respective secondary antibodies in 5% NGS/PBS/0.1% TritonX-100 for 2hrs in the dark at room temperature. The secondary antibodies used are listed in Supplementary Table. The spinal cords were then washed 3 times for 15mins with 0.1% TritonX-100/PBS(1X). The spinal cords were rinsed with PBS 1X. The spinal cords were then rinsed with 70% ethanol, and then incubated with Auto-fluorescent eliminator (EMD Millipore) for 5mins. The spinal cords were then rinsed 2 times for 5mins with 70% ethanol. Spinal cords were rehydrated in PBS (1X) for 5mins and then transferred to phosphate buffer (1X). Using a dissecting microscope, spinal cord sections were then mounted onto Superfrost Plus charged slides (Fisherbrand) and allowed to dry. 200µL of Fluoromount with DAPI (Sigma) was then added to the slide and a #1.5 22×48mm coverslip (Fisherbrand) was mounted on top. Confocal images were taken by an Olympus FV1000 Laser Scanning Confocal.

### RNA Extraction

Spinal cord and brain tissue was isolated as described above and then frozen at-80degC. Tissue samples were homogenized in 1ml of TRIZOL reagent with a power homogenizer (Fisher Scientific). The sample was then transferred to a phase lock gel tube, shaken vigorously, and incubated at room temperature for 5mins. 200µL of chloroform was added, shaken vigorously again for 15s and then incubated for 2mins at room temperature. The sample is then centrifuged for 15mins at 12,000gs at 4deg C. The aqueous phase is then mixed with an equal volume of ethanol and then the RNA is isolated using a PureLink RNA mini kit by manufacturer’s instructions.

### qPCR

RNA (200ng) was reverse transcribed (Superscript IV Reverse Transcriptase (Thermo Fisher)) and the output volume of 20µL was diluted in nuclease-free water to 40µL for a working concentration of 5ng/µL. Real time PCR was performed using SYBR Green PCR Master Mix (Applied Biosystems) on a QuantStudio 3 System (Applied Biosystems) with reaction specificity confirmed by melt curve analysis. For qPCR primer sequence, see Supplementary Table.

### Protein Tissue Extraction

Spinal cord and brain tissue was isolated as described above and then frozen at-80degC. Tissue samples were homogenized in 500ul of RIPA lysis buffer (Sigma) supplemented with protease inhibitors. The tissue was then passed through a 70uM cell filter where the lysate was collected in a 1.5mL microtube. The lysate was then agitated for 30mins at 4deg C and then centrifuged for 20mins at 12,000rpms at 4deg C. The supernatant was then transferred to a pre-chilled microtube and stored at −80deg C.

### RNA Seq of NEMF^R86S^ MEFs

Concentrated RNA was sent for bulk RNAseq to Azenta. In brief, sample quality control and determination of concentration was performed using TapeStation Analysis by Azenta, followed by library preparation and sequencing. Computational analysis included in their standard data analysis package was used for data interpretation.

### Image Analyses

Images were analyzed in bulk through Cell profiler. Z-projections were taken from each image by maximum intensity and then separated by fluorophore. Nuclear/Cytoplasmic (N/C) Ratios were taken by comparing area of nucleus (DAPI) and area of the cytoplasm (Phalloidin or brightfield). Nuclear and Cytoplasmic Intensities were standardized to area.

---

**Figure S1: *Nemf*^R86S^ Mice display nuclear loss of NEMF in the primary motor cortex and show a specific motor neuron defect in the spinal cord.**

A) Primary Motor Cortex was isolated from 21-day old Wild Type and *Nemf*^R86S^ mice. Neurons in the ventral horn were immunostained for the nucleus (DAPI, blue) and NEMF (red). B) Nuclear/Cytoplasmic ratios of NEMF in WT and Nemf^R86S^. Data analyzed by unpaired two-tailed t-test. C-D) Lumbar Spinal Cords were immunostained for the nucleus (DAPI, blue) and neurons (Nissl, red), and NEMF (C) and TDP-43 (D) (green). Arrows indicate pathological NEMF or TDP-43 Nissl-positive neurons. (**** p<0.0001)

**Figure S2: *Nemf*^R86S^ MEFs display cytoplasmic mis-localization of nucleoporins and a ‘leaky’ nuclear pore.** A-B) qPCR validation of *Ltn1* and *Nemf* siRNA knockdown (n=3). C) Outline of Nuclear Isolation and seeding onto coverslips. D) Immunostaining of Nuclei (DAPI, blue) and fluorescently-tagged Dextrans (20kD-FITC, 40kD Texas-Red, and 70kD Texas-Red. E) Quantification of integrated intensity of dextrans within the nucleus. (ns p>0.05, ** p<0.01)

**Figure S3: Negative Controls for NEMF and Importin-β PLA**

A) PLA of NEMF and Importin-β alone, or without primary antibodies (red). Nuclei labeled with DAPI (blue).

**Figure S4: *Nemf*^R86S^ MEFs show cytoplasmic mis-localization of nucleoporins and an increase in an amyloid-like phenotype.**

A) Quantification of WT and *Nemf*^R86S^ MEF cell size in square microns using phalloidin as cell maker. Data analyzed by unpaired two-tailed t-test. (n=999-1000) B) Growth curves of WT (n=3, r^2^=0.99) and Nemf^R86S^ (n=3, r^2^=0.98) over 48 hours at 15min intervals. Data was analyzed by a nonlinear regression for Malthusian growth. C) Immunofluorescent staining of Nup153, Nup50, and Nup98 (green). Nuclei labeled with DAPI (blue). D) Quantification of the percentage of cells with cytoplasmic puncta from Fig 4. Data was analyzed by two-way ANOVA with Šídák’s multiple comparisons test (n=3). E) Immunofluorescence Stain of amyloid in WT and *Nemf*^R86S^ MEFs by AmyloGlo. Nuclei labeled with Lamin A/C. F) Quantification of the Integrated Intensity of AmyloGlo. G) Stain-Free total protein of Sarkosyl Soluble and Insoluble TDP-43. Data analyzed by two-tailed t-test. (n=30-60) (ns p>0.05, **p<0.01, ****p<0.0001)

**Figure S5: Late onset *Nemf*^R487G^ spinal motor neurons show appearance of phospho-TDP-43 at 6- and 12-months**

A) Immunofluorescent staining of phospho-TDP-43 (red), ChAT (green), Nissl (white) in WT, and *Nemf*^R487G^ lumbar spinal cord motor neurons at 21 days, 6 months, and 12 months. Nuclei labeled with DAPI (blue).

**Figure S6: IPZ and IVM Cell Viability Curves and Gene Ontology Analyses** A-B) Cell Viability Curves for Importazole (A) and Ivermectin (B) treated MEFs. Data analyzed by one-way with Tukey’s multiple comparison test C-D) Gene Ontology Analysis of significantly dysregulated genes in Nemf^R86S^ and Importazole-treated samples. E-F) qPCR relative fold change of *Bax* and *Sorl1* in MEFs, spinal cord, brain in WT and *Nemf*^R86S^ mice. Data analyzed by two-way anova with Šídák’s multiple comparisons test. (n=3). (ns p>0.05, *p<0.05, **p<0.01, **** p<0.0001).

**Figure S7: RNA Quality is maintained high through all reads.**

A) RNA quality scores throughout Sanger Sequencing Position as presented by Azenta B) Tapestation Analysis comparing RNA integrity number for WT, IPZ-treated, and IVM-treated RNA samples. Data analyzed by one-way anova with Tukey’s multiple comparison test. (ns p>0.05) C) qPCR log2FoldChange plotted over RNA for MEFs (r^2^=.76,*p<0.05), Spinal Cord (SC) (r^2^=.89,**p<0.01), and Brain (r^2^=.05,*p>0.05). A simple Linear regression determined best-fit model.

**Figure S8: qPCR Validation and *STMN2* expression in HEK293 cells**

A) qPCR validation of *LTN1 (RNF160), NEMF* (*SDCCAG1*), *Importin-β* (*KPNB*), Importin-α (KPNA), or *Transportin-1* (TPNO1) siRNA knockdown in HEK293 cells (n=3). B) qPCR validation of *NEMF, LTN1,* and *Importin-β* siRNA knockdown in SK-N-MC cells (n=3). Data analyzed by two-way ANOVA with Šídák’s multiple comparisons test. C-D) qPCR relative fold change of *STMN2* or *APOE* treated with *LTN1, NEMF*, *Importin-β*, Importin-α, or *Transportin-1*siRNAs in HEK293 cells (n=3). E) qPCR relative fold change in DMSO Control and Importazole (IPZ) or Ivermectin (IVM) treated HEK293 for *STMN2* (n=3). Data analyzed by one-way anova with Tukey’s multiple comparison test (ns p>0.05, *p<0.05, **p<0.01, ***p<0.001 **** p<0.0001)

