## supplemental files for "Importin-β specific nuclear transport defects recapitulate phenotypic and transcriptional alterations seen in neurodegeneration"

**Supplementary Figures**

### Figure S1

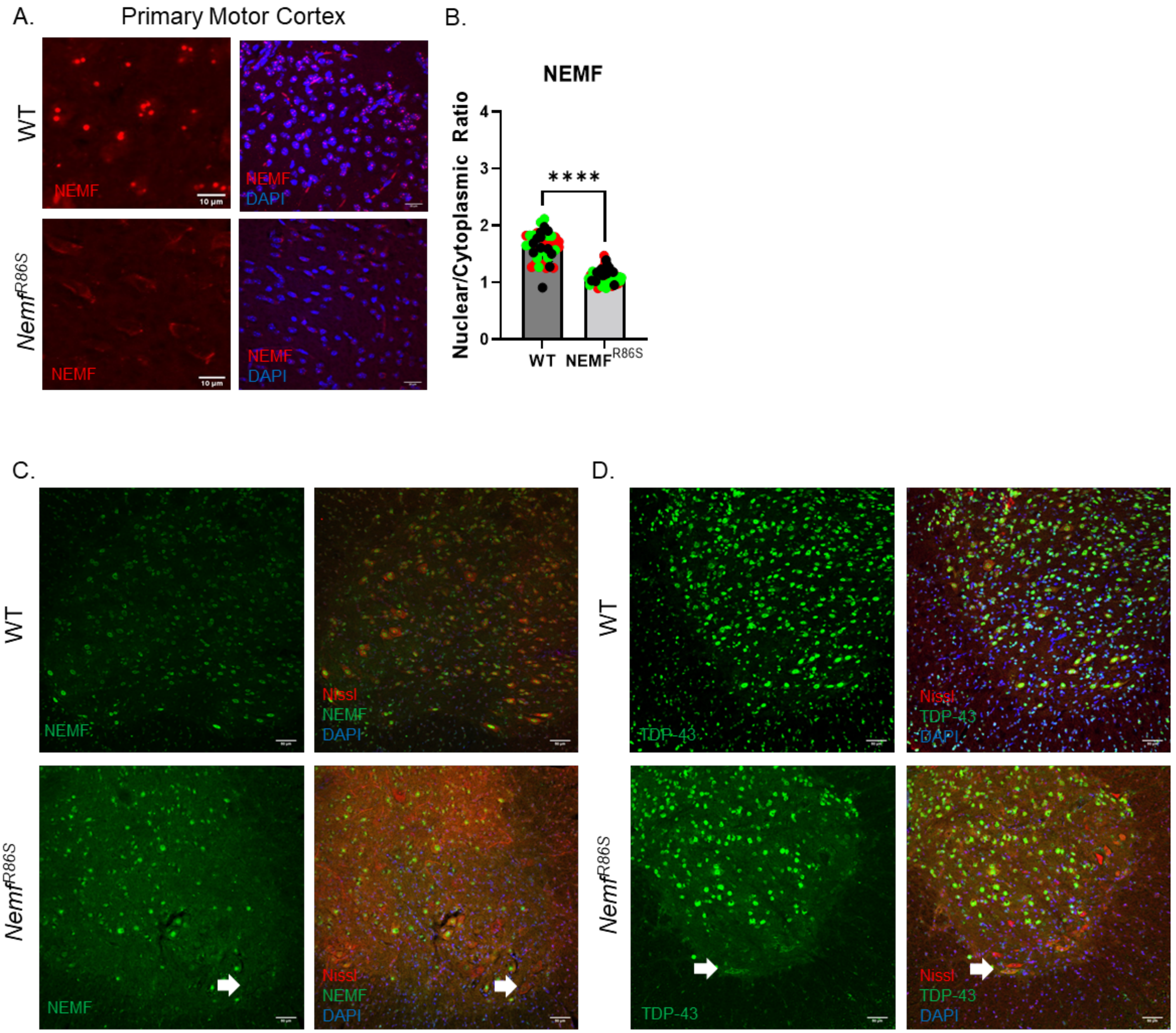

#### Figure S2

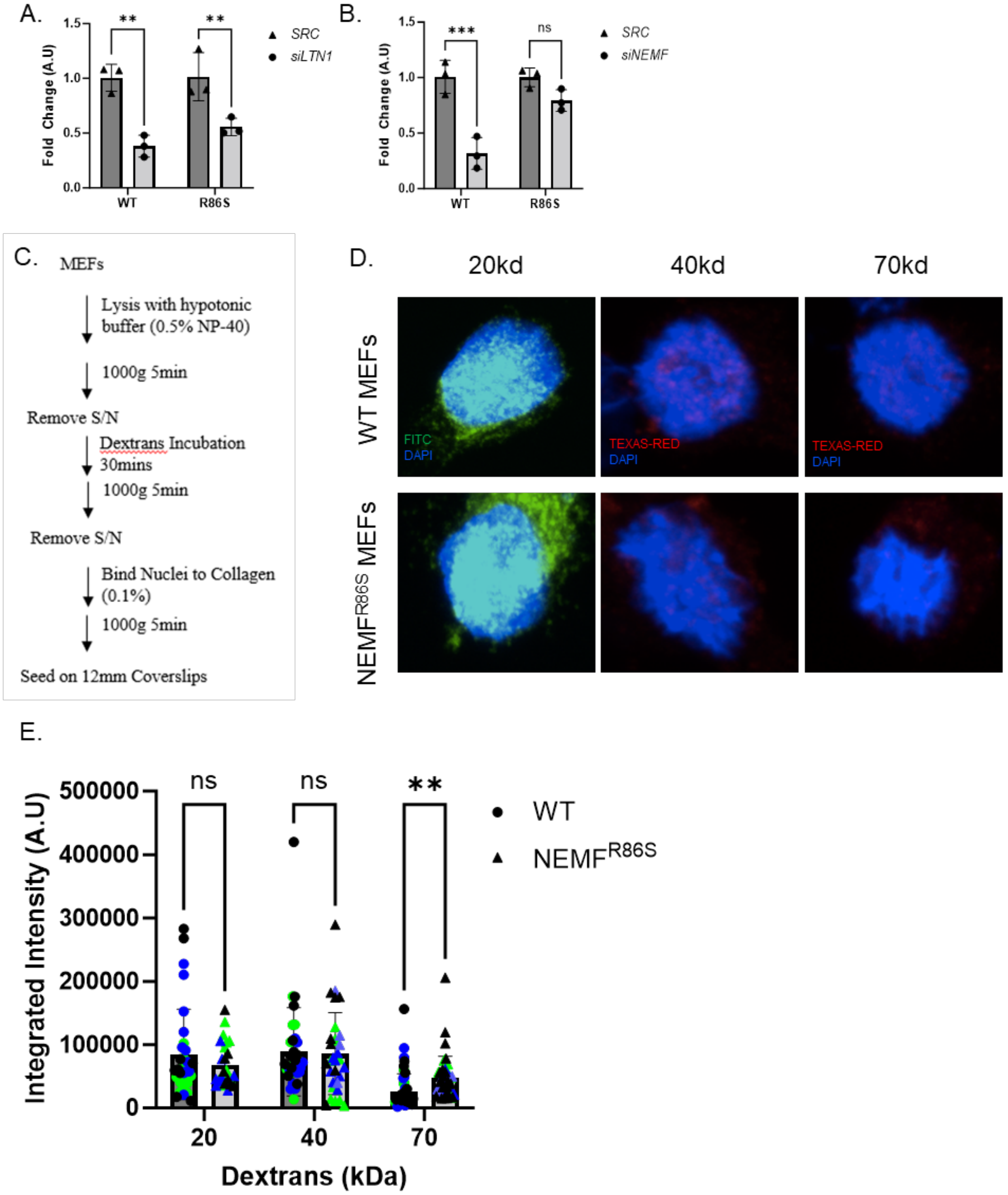


728

#### Figure S3

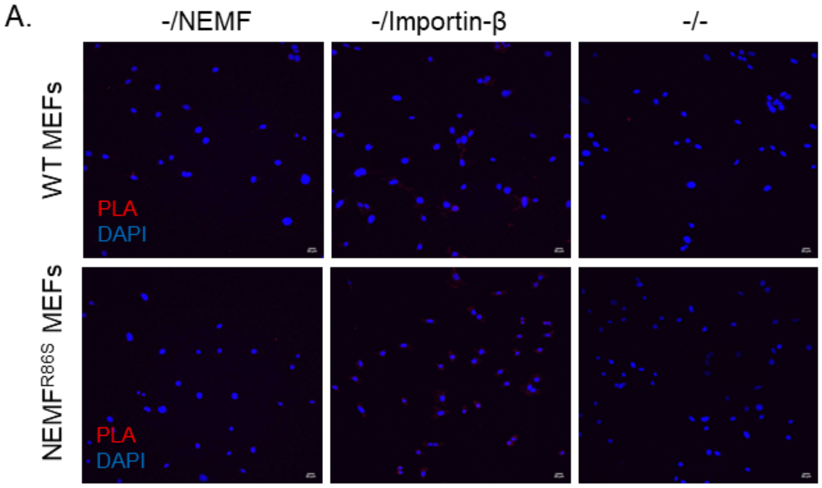


729

### Figure S4

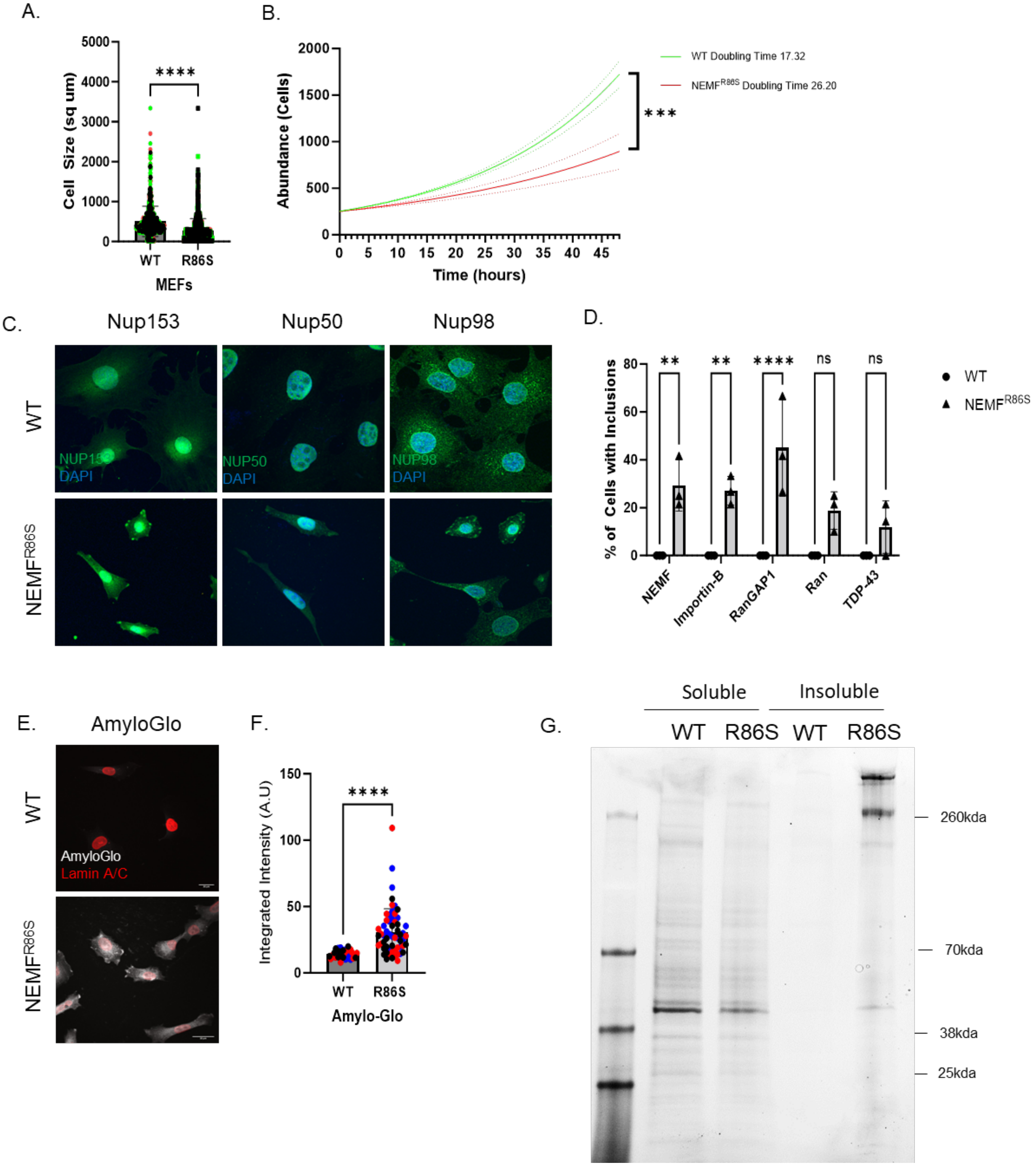
A.
5000
4000
3000
2000
1000
0
Cell Size (sq um)
****
WT
R86S
MEFs
B.
2000
1500
1000
500
0
Abundance (Cells)
0 5 10 15 20 25 30 35 40 45
Time (hours)
WT Doubling Time 17.32
NEMF$^{R86S}$ Doubling Time 26.20
***
C.
Nup153
Nup50
Nup98
WT
NEMF$^{R86S}$
NUP153
DAPI
NUP50
DAPI
NUP98
DAPI
D.
80
60
40
20
0
% of Cells with Inclusions
**
**
****
ns
ns
NEMF
Importin-B
RanGAP1
Ran
TDP-43
WT
NEMF$^{R86S}$
E.
AmyloGlo
WT
NEMF$^{R86S}$
AmyloGlo
Lamin A/C
F.
150
100
50
0
Integrated Intensity (A.U)
****
WT
R86S
Amylo-Glo
G.
Soluble
Insoluble
WT
R86S
WT
R86S
260kda
70kda
38kda
25kda

### Figure S5

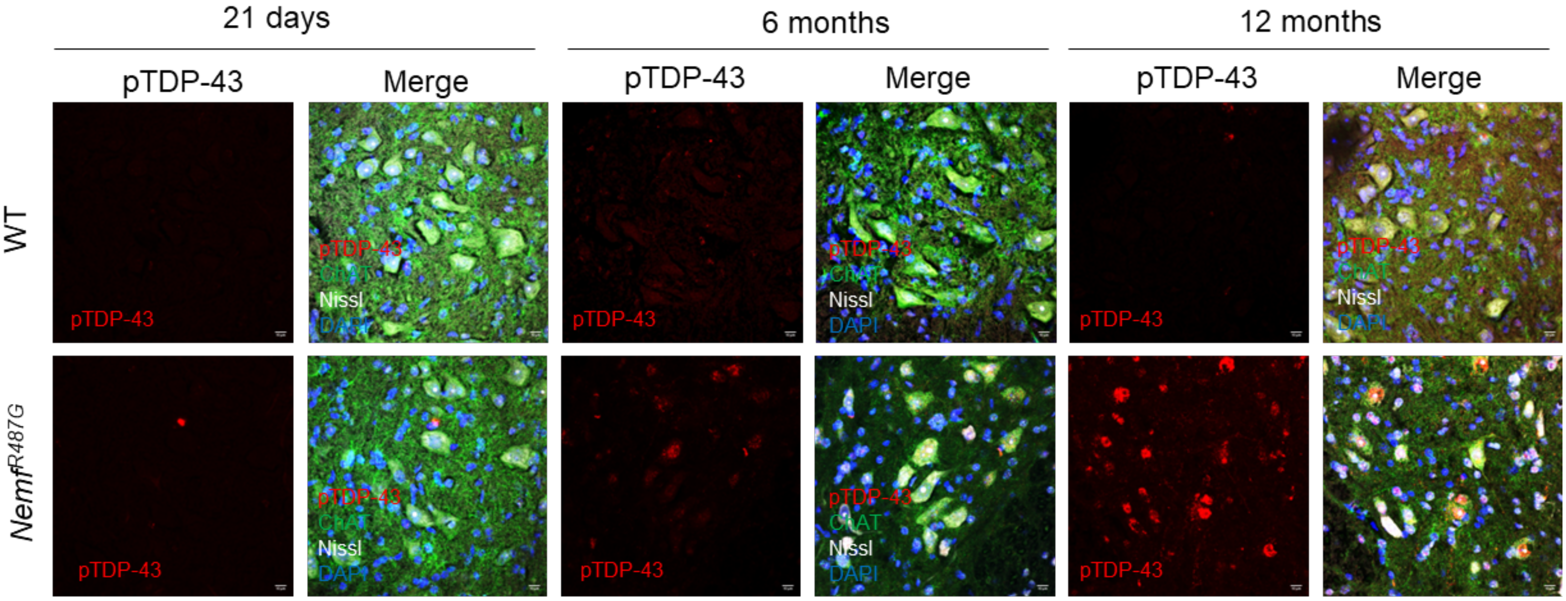
21 days
6 months
12 months
pTDP-43
Merge
WT
NemfR487G
pTDP-43
ChAT
Nissl
DAPI

### Figure S6

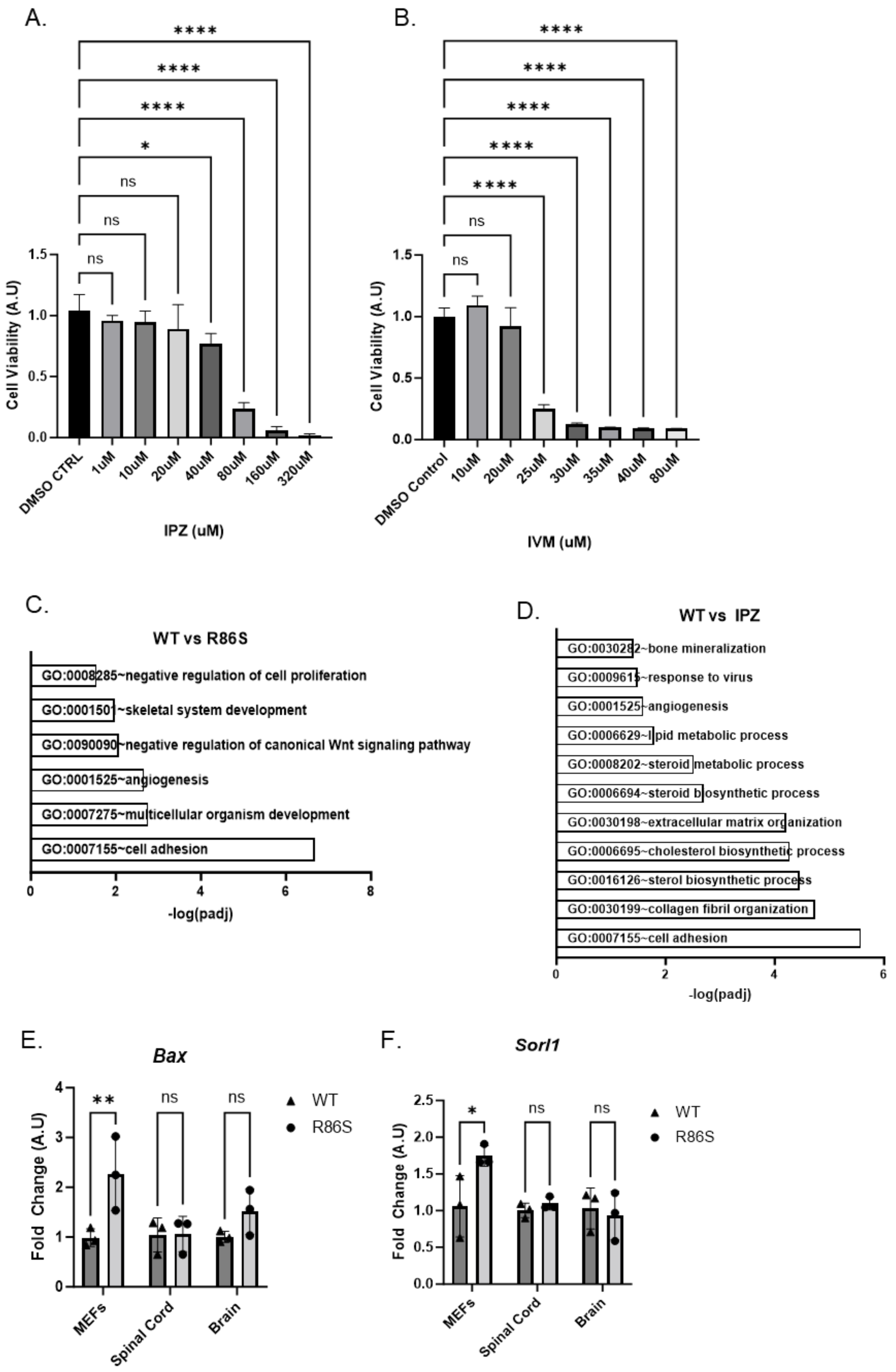
A.
Cell Viability (A.U)
IPZ (uM)
DMSO CTRL
1uM
10uM
20uM
40uM
80uM
160uM
320uM
ns
*
****
B.
Cell Viability (A.U)
IVM (uM)
DMSO Control
10uM
20uM
25uM
30uM
35uM
40uM
80uM
C.
WT vs R86S
GO:0008285~negative regulation of cell proliferation
GO:0001501~skeletal system development
GO:0090090~negative regulation of canonical Wnt signaling pathway
GO:0001525~angiogenesis
GO:0007275~multicellular organism development
GO:0007155~cell adhesion
-log(padj)
D.
WT vs IPZ
GO:0030282~bone mineralization
GO:0009615~response to virus
GO:0001525~angiogenesis
GO:0006629~lipid metabolic process
GO:0008202~steroid metabolic process
GO:0006694~steroid biosynthetic process
GO:0030198~extracellular matrix organization
GO:0006695~cholesterol biosynthetic process
GO:0016126~sterol biosynthetic process
GO:0030199~collagen fibril organization
GO:0007155~cell adhesion
-log(padj)
E.
Bax
Fold Change (A.U)
**
WT
R86S
MEFs
Spinal Cord
Brain
F.
Sorl1
Fold Change (A.U)

#### Figure S7

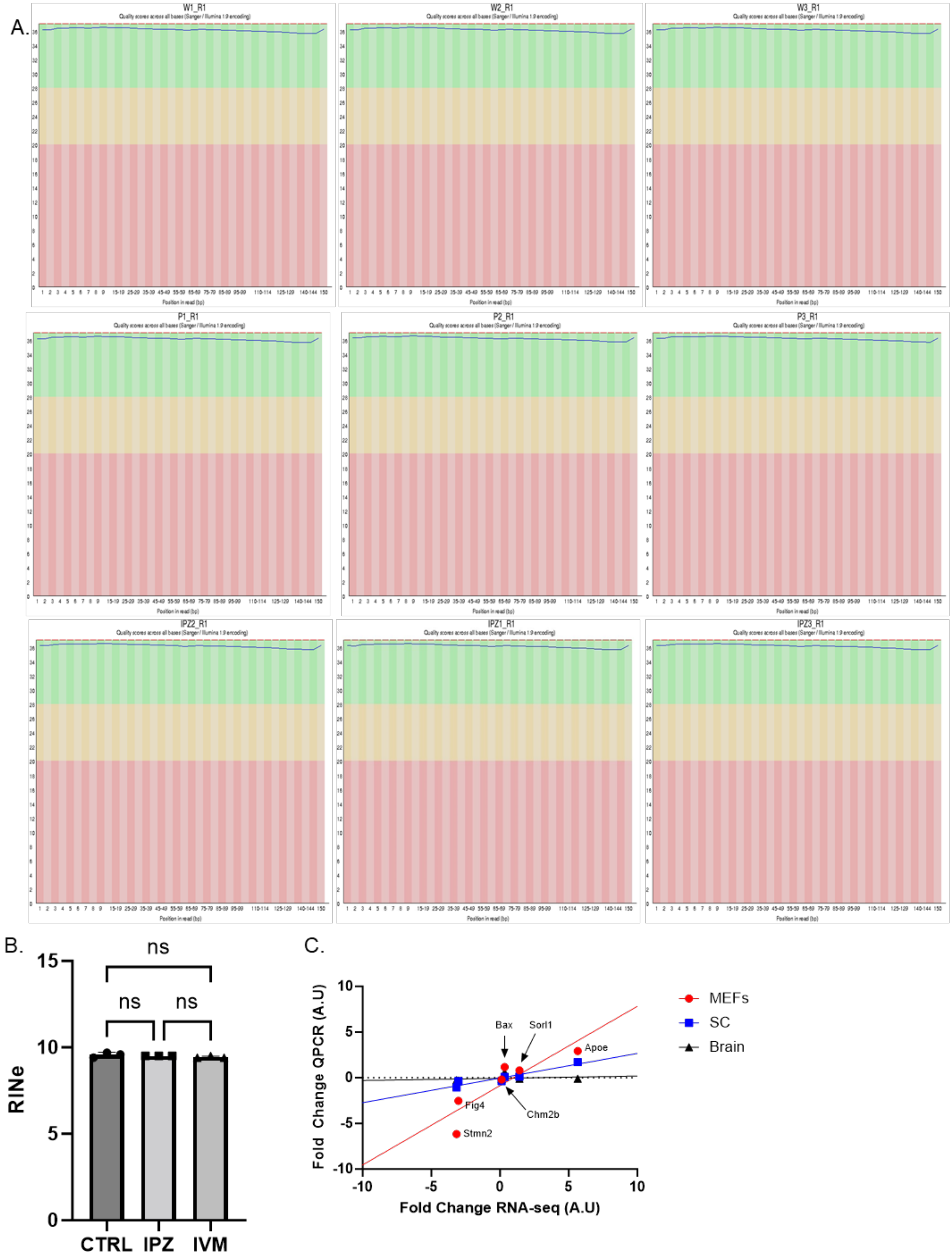

A.
W1_R1
W2_R1
W3_R1
P1_R1
P2_R1
P3_R1
IPZ2_R1
IPZ1_R1
IPZ3_R1
B.
ns
ns
ns
RINe
15
10
5
0
CTRL
IPZ
IVM
C.
Fold Change QPCR (A.U)
Fold Change RNA-seq (A.U)
10
5
0
-5
-10
Bax
Sorl1
Apoe
Fig4
Chm2b
Stmn2
MEFs
SC
Brain

### Figure S8

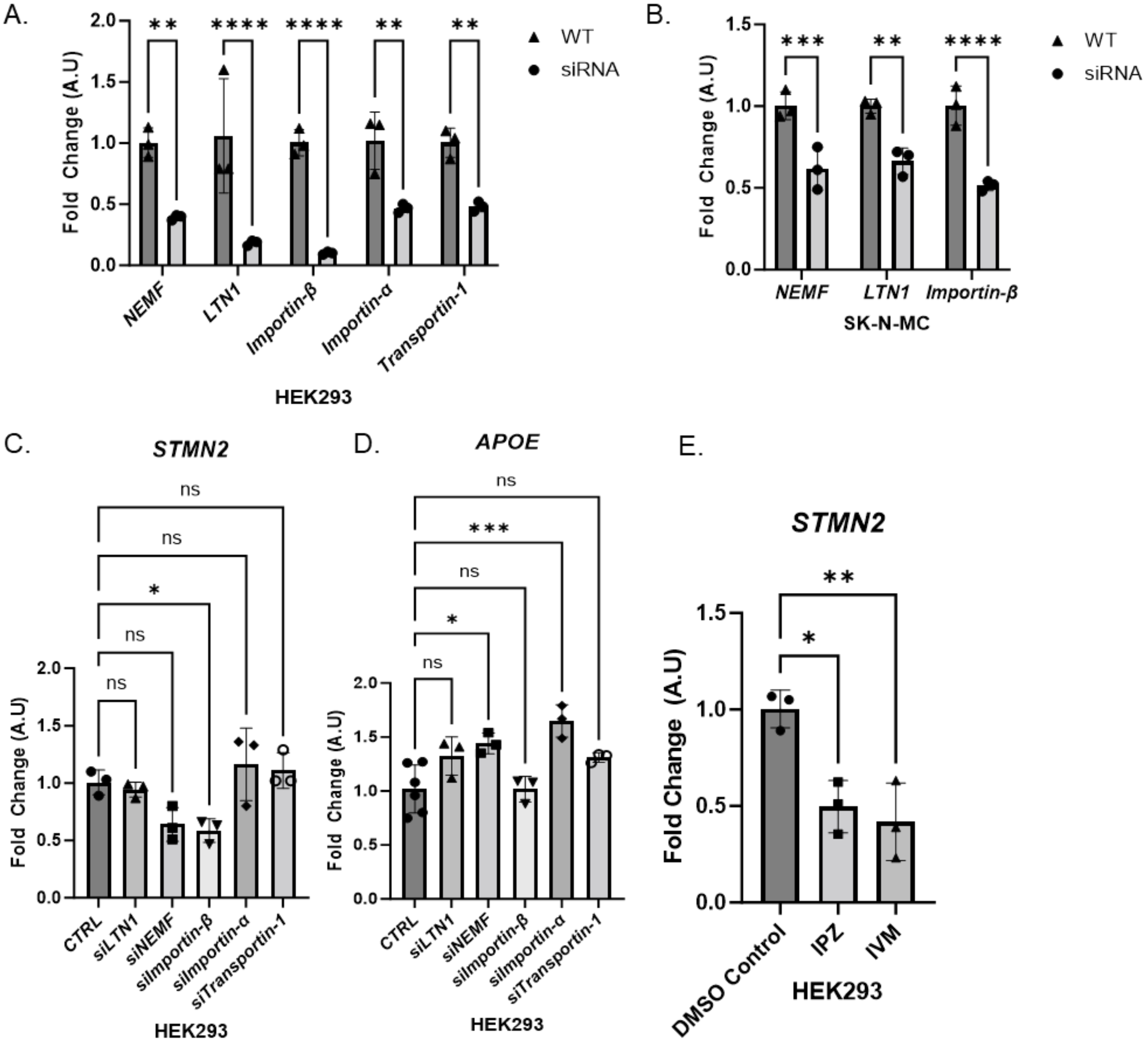
A.
Fold Change (A.U)
** **** **** ** **
NEMF LTN1 Importin-β Importin-α Transportin-1
HEK293
WT
siRNA
B.
Fold Change (A.U)
*** ** ****
NEMF LTN1 Importin-β
SK-N-MC
WT
siRNA
C.
STMN2
ns ns * ns ns
Fold Change (A.U)
CTRL siLTN1 siNEMF siImportin-β siImportin-α siTransportin-1
HEK293
D.
APOE
ns *** ns * ns
Fold Change (A.U)
CTRL siLTN1 siNEMF siImportin-β siImportin-α siTransportin-1
HEK293
E.
STMN2
** *
Fold Change (A.U)
DMSO Control IPZ IVM
HEK293

### Supplementary Tables

| Antibody | Company | Cat # |
|---|---|---|
| Alexa Fluor 594 g@ms IgG2a | Invitrogen | A21135 |
| Alexa Fluor 594 g@Rb Ig | Invitrogen | A11034 |
| Alexa Fluor 488 g@Chk IgY | Invitrogen | A11039 |
| Alexa Fluor 594 g@Chk IgY | Invitrogen | A11042 |
| Alexa Fluor 594 g@ms IgG1 | Invitrogen | A21125 |
| Alexa Fluor 594 g@Rb Ig | Invitrogen | A11037 |
| Alexa Fluor 594 g@ms IgG2b | Invitrogen | A21141 |
| IRDye 800CW Goat anti-Mouse IgG1 | Licor | 926-32350 |
| IRDye 800CW Goat anti-Mouse IgG2a | Licor | 926-32351 |
| IRDye 800CW Goat anti-Rabbit IgG | Licor | 926-32211 |
| IRDye 800CW Donkey anti-Chicken IgG | Licor | 926-32218 |

| Antibody | Company | Cat # |
|---|---|---|
| NEMF | mybiosource | MBS821574 |
| TDP43 | proteintech | 10782-2-AP |
| MAP2 | proteintech | 17490-1-AP |
| RPL3 | proteintech | 66130-1-Ig |
| Listerin | Abcam | ab104375 |
| Lamin A/C | Cell Signaling | 4C11 |
| Importin-B | proteintech | 10077-1-AP |
| Importin-B | Invitrogen | MA3-070 |
| RanGAP1 | Invitrogen | 33-0800 |
| ChAT | Aves | CAT |
| GFP | Aves | GFP-1020 |
| Nup153 | proteintech | 14189-1-AP |
| Nup98 | Cell Signaling | C39A3 |
| Nup50 | Invitrogen | PA5-28452 |
| Ran | proteintech | 10469-1-AP |
| RPS6 | Santa Cruz | sc-74459 |
| Phalloidin | Invitrogen | A12381 |
| mCherry | Abcam | ab125096 |
| TIAR | Cell Signaling | 5137 |
| pTDP-43 (h) | Protein-Tech | 10782-2-AP |
| pTDP-43 (m) | Kind Gift of Dr. Leonard Petrucelli, Mayo Clinic | |

| Primer | Forward | Reverse |
|---|---|---|
| *Apoe (mouse)* | GACCCAGCAAATACGCCTG | CATGTCTTCCACTATTGGCTCG |
| *Gapdh (mouse)* | GGCAAATTCAACGGCACAGT | GGGTCTCGCTCCTGGAAGAT |
| *Stmn2 (mouse)* | TGTCACTGATCTGCTCCTGC | TGGGAGATGGTGGCTTCAAG |
| *Bax (mouse)* | CTACAGGGTTTCATCCAG | CCAGTTCATCTCCAATTCG |
| *Chmp2b (mouse)* | GGACCGAGCAGCCTTAGAG | CCAATCTTGGCCATCTTCTTA |
| *Sddcag1 (human)* | GACATGGAGACACTGGCAAGTTG | CCTCTTGGTCAACCTGAGTCATG |
| *Rnf160 (human)* | GATGAGGCAGTCTCTTCCTATGC | GCTGAAGTACACAAATGGCTGGG |
| *Kpnb1 (human)* | CTGCTTCCTGAAGCTGCCATCA | CTTCAGCCAGACTGGAGAAAGC |
| *Kpna1 (human)* | TTCCAAAAGCCCAGAGCAACAGC | CCACTACTCCTGGTGTGCTGAT |
| *Tpno1 (human)* | TGGCTGAAGGACTTGGAGGCAA | TGTGAGGTCACCTAACAGGGCA |
| *Nemf (mouse)* | GCTGCACAAGTGAACTCATGGC | GAGTTCTACGTCTCCAGGCATTC |
| *Ltn1 (mouse)* | TGTTCTGTGGCTGAAGGACCAG | CAAGAAGAGACGTGTCCTCACTC |
| *Gapdh (human)* | GTCTCCTCTGACTTCAACAGCG | ACCACCCTGTTGCTGTAGCCAA |
| *18S (rRNA)* | CTCAACACGGGAAACCTCAC | CGCTCCACCAACTAAGAACG |
| *28S (rRNA)* | CTAAATACCGGCACGAGACC | TTCACGCCCTCTTGAACTCT |
| NMNAT2 (exon 1-2) | CAGTGCGAGAGACCTCATCCC | ACACATGATGAGACGGTGCCG |
| NMANT2 (exon 4-5) | GAGTGCTATCAGGACACCTGG | GTGGGCACATTGCTGTTCTGG |

### Supplementary Tables

| Plasmid | Company |
|---|---|
| pCMV6-AC-IRES-GFP | AddGene |
| pCMV6-AC-IRES-GFP-3X-NLS | This paper |
| pmCherry-FUS (PY)-NLS | This paper |
| pREV-FUS-NLS-PKINES-mCherry | This paper |
| SiRNA | Company |
| SRC (mouse) | Ambion |
| NEMF (mouse) | Ambion |
| LTN1 (mouse) | Ambion |
| Sddcag1 (human) | Thermo |
| RNF160 (human) | Thermo |
| Kpnb1 (human) | Thermo |
| Kpna1 (human) | Thermo |
| Tpno1 (human) | Thermo |

| Cell Culture | Company | Cat # |
|---|---|---|
| Dulbecoo's Modified Eagle's Medium | Corning | 10-017-CV |
| Eagle's Minimum Essential Medium | ATCC | 30-2003 |
| FBS 1X | Gibco | 26140079 |
| Penstrep 100X | Gibco | 15140122 |
| Glutamax 100X | Gibco | 35050061 |
| Non-essential Amino-Acids 100X | Gibco | 11140050 |
| Trypsin-EDTA (0.05%) | Gibco | 25300062 |

| **Product** | **Company** | **Catalog #** |
|---|---|---|
| 12mm Coverslips | Carolina Biological Supply | 633029 |
| gelatin | Millipore Sigma | G6650 |
| Lipofectamine 3000 | Life Technologies | L3000001 |
| Lipofectamine 2000 | Life Technologies | 11668500 |
| Importazole | Sigma | SML0341 |
| Ivermectin | Sigma | PHR1380 |
| Prolong Glass with NucBlue | Life Technologies Corporation | P36983 |
| RIPA lysis buffer | Sigma Aldrich | R0278 |
| Pierce BCA | Thermo Scientific | 23227 |
| dTT | Bio-rad | 1610611 |
| 4X laemli buffer | Bio-Rad | 1610747 |
| Pre-Cast Gel (4-15% MP TGX Gel 10W 50 ul) | Bio-Rad | 4561084 |
| Tris/Glycine/SDS 10X | Bio-Rad | 1610732 |
| Chameleon 800 MW ladder | LI-COR | 928-80000 |
| Tris/Glycine Buffer 10X | Bio-Rad | 1610771 |
| Auto-fluorescent eliminator | EMD Millipore | 2160 |
| Superfrost Plus | Fisherbrand | 12-550-14 |
| Fluoromount with DAPI | Southern Biotech | 0100-20 |
| 22x48mm coverslip | Fisherbrand | 12-550-14 |
| Superscript IV Reverse Transcriptase | Thermo Fisher | 18090010 |
| SYBR Green PCR Master Mix | Thermo Fisher | 4472918 |
